# ThunderBolt: An interactive data sharing and analysis platform for large-omics experiments

**DOI:** 10.1101/2024.07.24.605014

**Authors:** Thomas Andrew Geddes, Rima Chaudhuri, Benjamin L. Parker, Pengyi Yang, James G. Burchfield

## Abstract

**Summary:** Mass-spectrometry (MS) datasets present a unique set of challenges that make in-depth bioinformatics analysis non-trivial, with analysis requiring both expertise and time. Often these datasets have unique structures that need to be dealt with on an individual basis. Currently, tools providing a fast, interactive and guided way of exploring and analysing these data sets are not readily available. To this end, we have developed ThunderBolt: a highly interactive, point-and-click web-based application providing both bioinformaticians and biologists with a platform for i) searching and comparing multiple omics datasets, ii) fast data exploration and quality control, iii) interactive visualization, iv) pre-processing, v) statistical analysis and vi) functional and network enrichment analysis of large proteomics datasets using the Shiny framework.

**Availability:** ThunderBolt is a shiny-application accessible at https://thunderbolt.sydney.edu.au/

**Supplementary information:** 

## 1 Introduction

High throughput technologies such as Mass Spectrometry (MS) have become central to many facets of biology including proteomics, metabolomics and lipidomics. In the field or proteomics, MS is used to quantify the proteome of cells or tissues, detect post-translational modifications (PTMs) and identify novel protein-protein interactions (PPIs). These experiments produce large datasets. For example, one phosphoproteomics study identified nearly 40,000 phosphorylation sites on more than 5,000 proteins (Humphrey et al. 2013). The unique complexity of these data such as the large proportions of missing values that occur non-randomly mandate the use of MS-specific computational approaches. Identifying the structure of these missing values influences the choice of i) imputation method, ii) filtering options and iii) normalization method, and the order they are applied to the dataset.

Tools for fast, interactive searching, comparison across multiple omics datasets and for performing interactive quality control are not readily available. While established software such as MultiExperiment Viewer (MeV) (TM4 2005) from the TM4 microarray suite and Perseus (Tyanova et al. 2016) (for data analysis) and Cytoscape (Shannon et al. 2003) or GePhi (Bastian et al. 2009) (for network visualization) are available, the time and expertise needed by users to use these platforms is non-trivial. In particular, current software does not provide an interactive and customizable environment that can be easily adapted to the emerging needs of the field and new methods as they become available. To address this we have developed ThunderBolt, a Shiny application (Chang et al. 2015), to provide a well-automated, highly interactive, comprehensive set of tools for data sharing and MS-based proteomics and interactomics data analysis for experts and non-experts alike.

## 2 Results and Overview

ThunderBolt enables interactive analysis, visualization and sharing of large proteomics datasets using an intuitive easy-to-use point-and-click web interface. The Shiny platform on which ThunderBolt was built is highly adaptable: new modules can be included as required, and new R packages may be easily integrated as they become available. Such interactivity and flexibility are lacking in existing tools. Figure 1 summarizes the functionality of ThunderBolt.

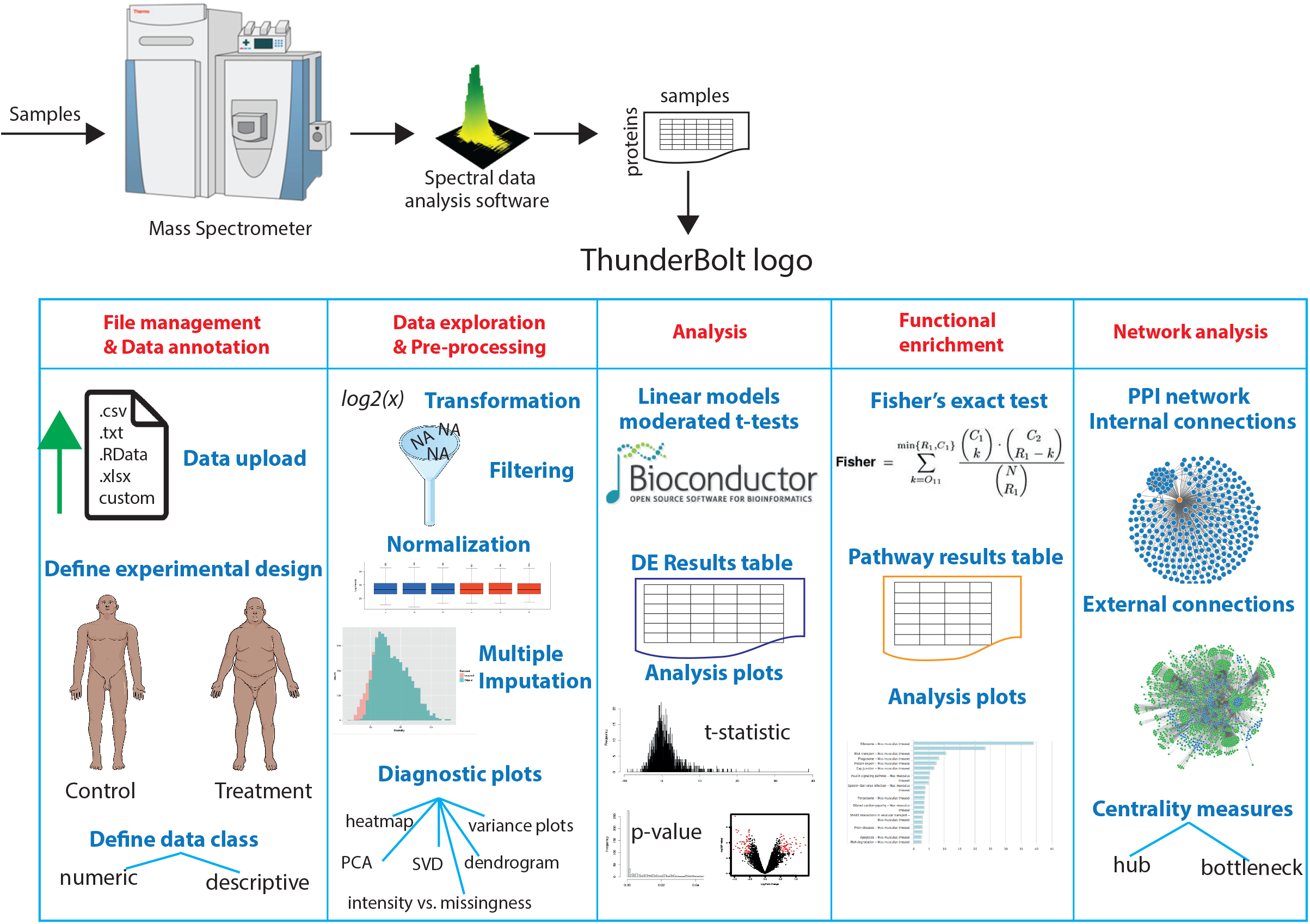

ThunderBolt contains eight different modules (Fig. 1). We have included an in-depth tutorial to demonstrate the usage and utility of these modules while exploring an interactomics dataset for the protein AKAP1. The functionality of these modules is summarized below:

### 2.1 Modules

#### File management and annotation

Users can upload their data in various file formats (RData, excel workbooks with multiple sheets, csv, tsv and custom delimited text files) and preview the data via a field-searchable table. Users may opt to save the file to the server to later restore and use in other modules or may proceed directly to the DE analysis pipeline without saving.

#### Data annotation

Once a dataset has been loaded into the DE pipeline, the user can annotate the data type and content of the columns to be included. The user defines the experimental design (controls versus treatments, IP antibodies etc.), to include descriptive and numerical columns, allowing ThunderBolt to group data for analysis.

#### Data exploration and pre-processing

Once the experimental conditions for the samples are defined, users can browse through a range of exploratory and diagnostic plots for quality control. Diagnostic plots include searchable protein expression, sample boxplots, proportion of missing values, sample and mean group variance, clustering, principal component analysis and eigenMS singular value decomposition (Karpievitch et al. 2012). A heatmap illustrates the global distribution of intensity ranks and missing values, separated by sample; another plot reflects the relationship between missingness and intensity by group. Thunderbolt facilitates a variety of pre-processing steps (see supplementary tutorial), the impact of which is immediately reflected in these diagnostic plots; this provides a global understanding of the dataset and guides subsequent pre-processing decisions.

#### Differential expression analysis

Con**Error! Bookmark not defined**.trol and experimental groups in the data can then be modelled to identify differentially expressed proteins using moderated t-tests (Smyth 2004). DE analysis results for each protein, including fold change (log scale); moderated t-statistic; raw and Benjamini-Hochberg FDR adjusted p-values, are tabulated and visualised interactively. The module permits selective filtering and subsequent download of these results based on specified fold change and raw-adjusted p-value cut-offs.

#### Functional enrichment

This module performs pathway over-representation analysis based on Fisher’s exact test. It accepts a list of proteins, either generated by the DE analysis module based on significance criteria or uploaded directly into the shiny app. ThunderBolt accepts a vector of Gene Symbols, UniprotID or EntrezID (for human and mouse) and performs a series of enrichment tests against pathway protein lists obtained from the KEGG pathways database (Kanehisa & Goto 2000), obtained via the KEGGREST library (Tenenbaum, 2021).. Significant enriched pathways from this analysis are listed in a table with additional visualisation. Enriched pathways can be saved to the server and accessed from the *Compare* module.

#### Network analytics

Similar to the functional enrichment module, DE proteins from the analysis module can be passed directly into this module for network generation based on known experimental PPIs from the STRING database (Snel et al. 2000). Users have the option to obtain a network including direct links between the input proteins (internal) or all proteins interacting with at least two members of the protein list (external) to generate a more complex and expansive interaction plot. ThunderBolt calculates centrality measures of the network including hub centrality score and betweenness, which can be downloaded as tables. Interactive networks are visualized using D3.js (Bostock et al. 2011); they can be exported in other graph formats (such as graphML or DOT) for visualization and editing purposes using other software tools.

#### Search

Uploaded and annotated datasets stored to the server can be easily accessed online by collaborators. Users can search by expression changes across differentially expressed (DE) proteins or specific protein(s) across multiple datasets/experiments and view or download resulting subsets.

#### Compare

This is a data linkage platform where users can compare and contrast multiple protein lists, biological pathways or other descriptive terms obtained from datasets/analyses, revealing patterns and similarities across experiments. Comparison overlaps can be visualized through Venn diagrams, tables or heatmaps.

## Conclusions

We have presented an interactive web-platform, ThunderBolt. This is an intuitive, interactive point and click web-interface for data sharing, exploration, fast interactive pre-processing, data analysis and visualization of large -omics datasets.

## Supporting information

Supplemental Data_Tutorial

## Funding

This work is supported by ARC/DP (DP170100654) to JYHY and PY; NHMRC/ECF (1072129) to BLP; ARC/DECRA (DE170100759) to PY; and NHMRC/CRE (1135285) to JYHY., ARC/DP (DP180103482) to DEJ and JGB and an NHMRC /Investigator Grant (1173469) to P.Y

## Conflict of Interest

none declared.

## Notes

### Competing Interest Statement

The authors have declared no competing interest.

