## Supplemental Data_Tutorial for "ThunderBolt: An interactive data sharing and analysis platform for large-omics experiments"

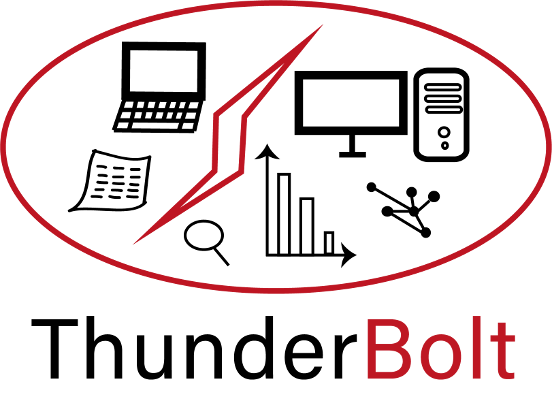

**Tutorial**

Created by:

Thomas Andrew Geddes

Rima Chaudhuri, Ph.D

at

Charles Perkins Centre

University of Sydney

Last updated:

22 October 2021

***Note:*** *On the thunderbolt.sydney.edu.au website, any datasets saved to the server are publicly accessible, so caution is advised. Datasets uploaded to this website may be periodically deleted****. Saving data is not required to use DE Analysis Pipeline, Pathway Analysis, Network Analysis Modules and Compare***

### Access

Thunderbolt is a web based application developed in Shiny that is designed to provide a robust pipeline for the analysis of MS based proteomics data. The application can be accessed from [thunderbolt.sydney.edu.au](http://www.thunderbolt.sydney.edu.au).

### Module: FILE MANAGEMENT

Files can be uploaded to the server and handled in three different ways as shown in Fig. T1-A; the associated tooltip provides a brief explanation.

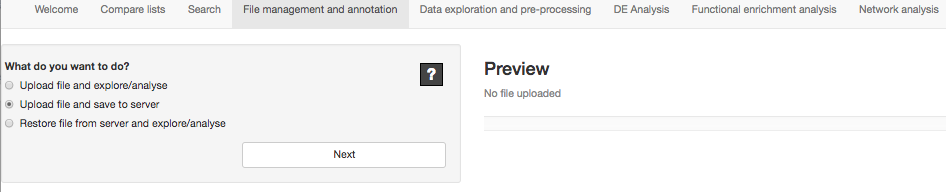

Fig. T1-A

Option #1: “Upload file and explore/analyse”. This option allows users to upload any unprocessed data (e.g. intensities from MaxQuant) and explore and perform DE analysis without retaining any history within Thunderbolt. In case of an unfortunate system crash, no data will be saved in the server and the user has to start the upload/analysis from scratch.

Option #2: “Upload file and save to server”. This allows users to upload data and save it to the server, adding any appropriate details (prompted after clicking the “next” button). This allows for data recovery in case of a crash. In Fig. T1-B, we use this option to upload a sample interactomics dataset with 1000 records (this can be loaded from the “DE Pipeline” → “Getting started” page in Thunderbolt). Fig. T1-C shows the table displayed once upload is completed. Once the dataset has been uploaded and correctly displayed as a table, several details can be added to name the dataset (Fig. T1-D). Using its assigned short name to select the dataset, it can be consequently searched or used for further analysis.

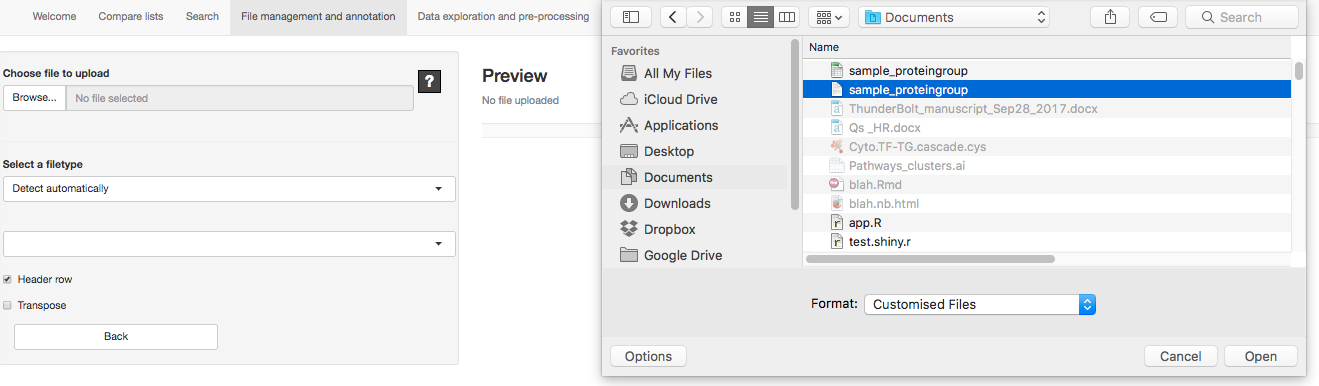

Fig. T1-B

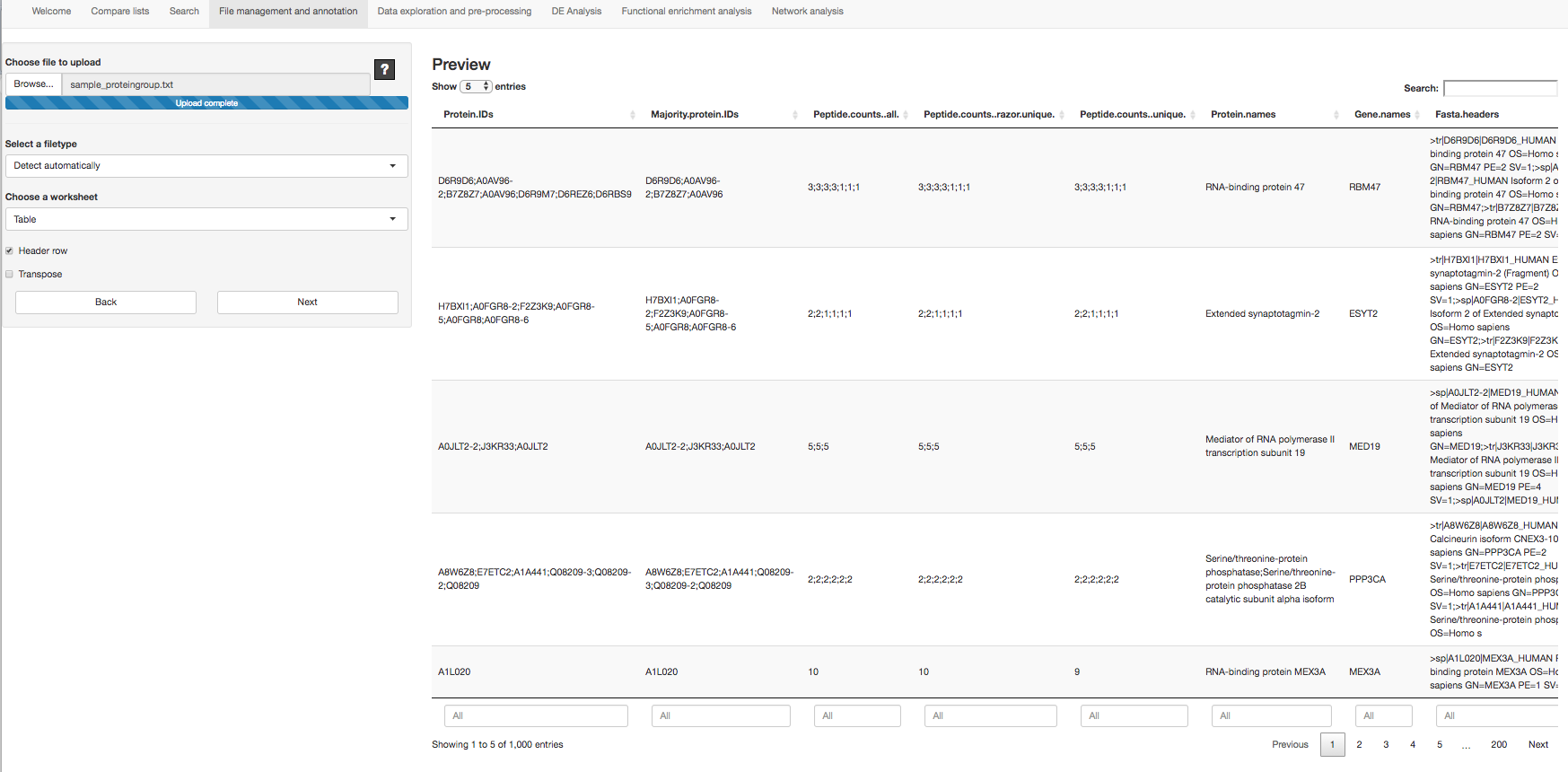

Fig. T1-C

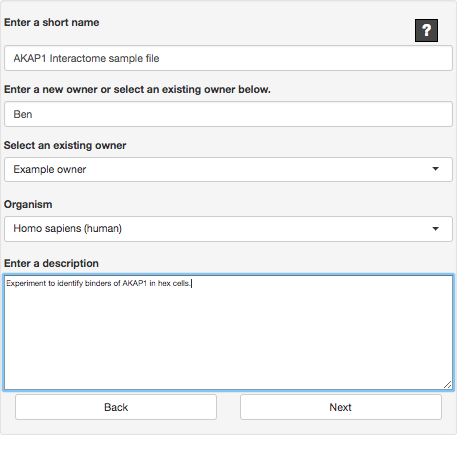

Fig. T1-D

Option #3: “Restore file from server and explore/analyse”. This option allows users to open a previously saved dataset and begin exploring it. Option #3 is usable only if a dataset has been uploaded using Option #2.

### Module: ANNOTATION

Once uploaded or restored, a dataset can be annotated with details of the experimental design. A brief explanation is provided on the page as shown in Fig. T2-A.

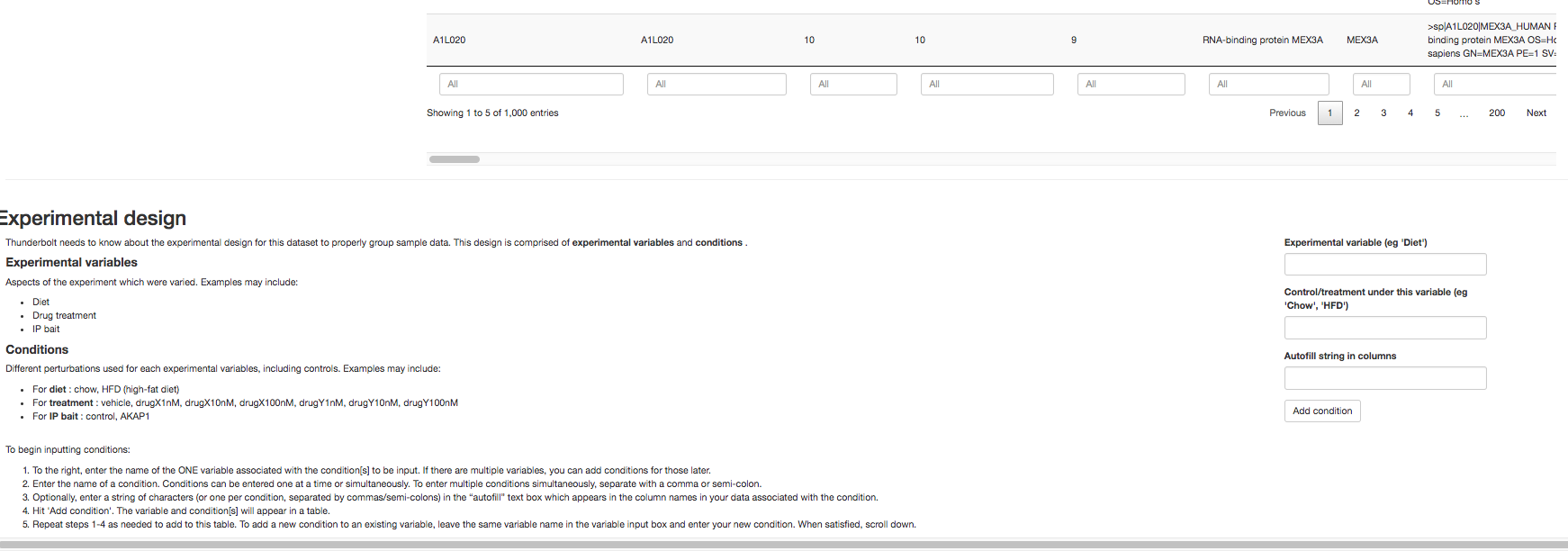

Fig. T2-A

In our example AKAP1 interactomics dataset, we have the following 4 experimental groups, n = 3, each containing 3 replicates. Experimental variables for this dataset include the bait for the pull-down assay, including the empty vector and AKAP1 conditions, and stimulus condition including basal and stimulated (“AA”). Combining these variables results in the four conditions: “EV_B”, “EV_AA”, “WT_B” and “WT_AA”.

As per the instructions in T2-A, we enter conditions for one experimental variable at a time. We first enter the conditions for the variable “Bait”: “Control” and “AKAP1”, as shown in Fig. T2-B.

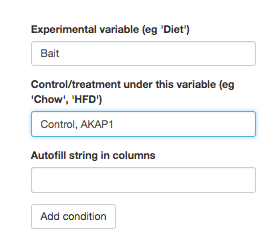

Fig. T2-B

Upon clicking “Add condition” the design table is updated (Fig. T2-C) and we can proceed to add the second variable “Stimulus” and its two associated conditions, “Basal” and “AA”. Once this is complete, we may annotate each of the columns in the dataset as demonstrated in Fig. T2-D. The metadata or descriptive columns that we want to retain through the analysis process can be selected and made distinct from numeric data columns at this step. If a specific string is common to the column names of all numeric columns of interest, such as “LFQ.intensity”, these columns can be selected by populating the field “Mark all columns as numeric containing string”. This designates as numeric any column whose header contains the string “LFQ.intensity”.

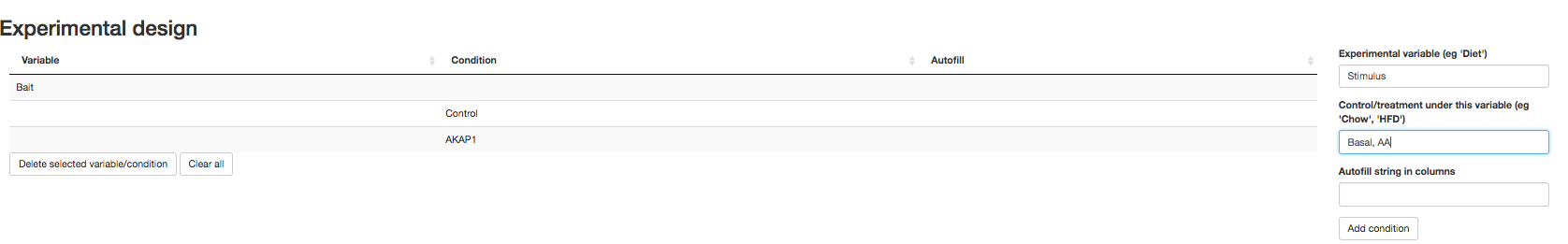

Fig. T2-C

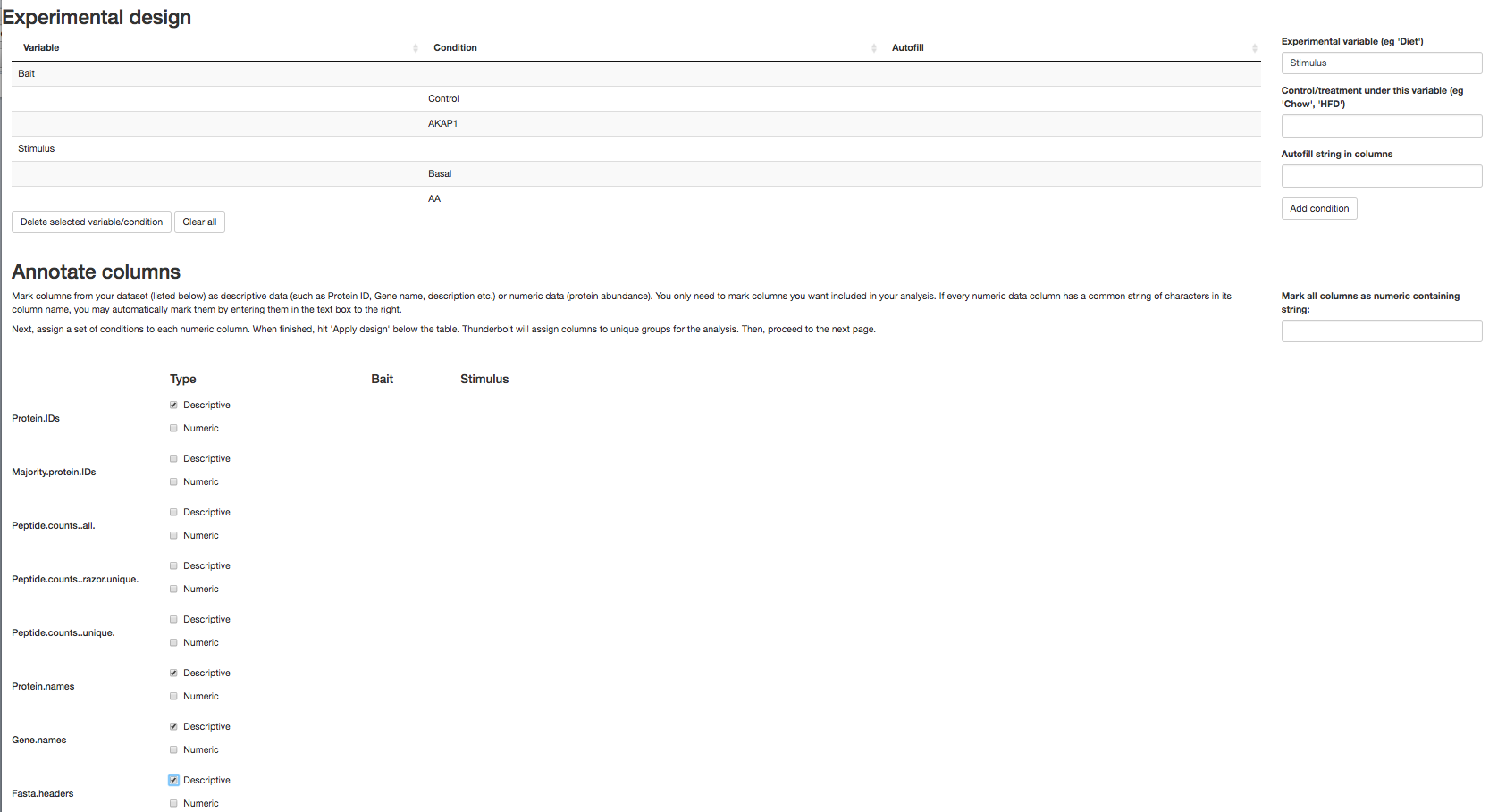

Fig. T2-D

The numeric columns needs to be further annotated by the user to assign which experimental group it belongs to using the drop-down boxes as shown in Fig. T2-E. If the user filled in the “Autofill string in columns” field when inputting experimental variables, Thunderbolt attempts to set these automatically.

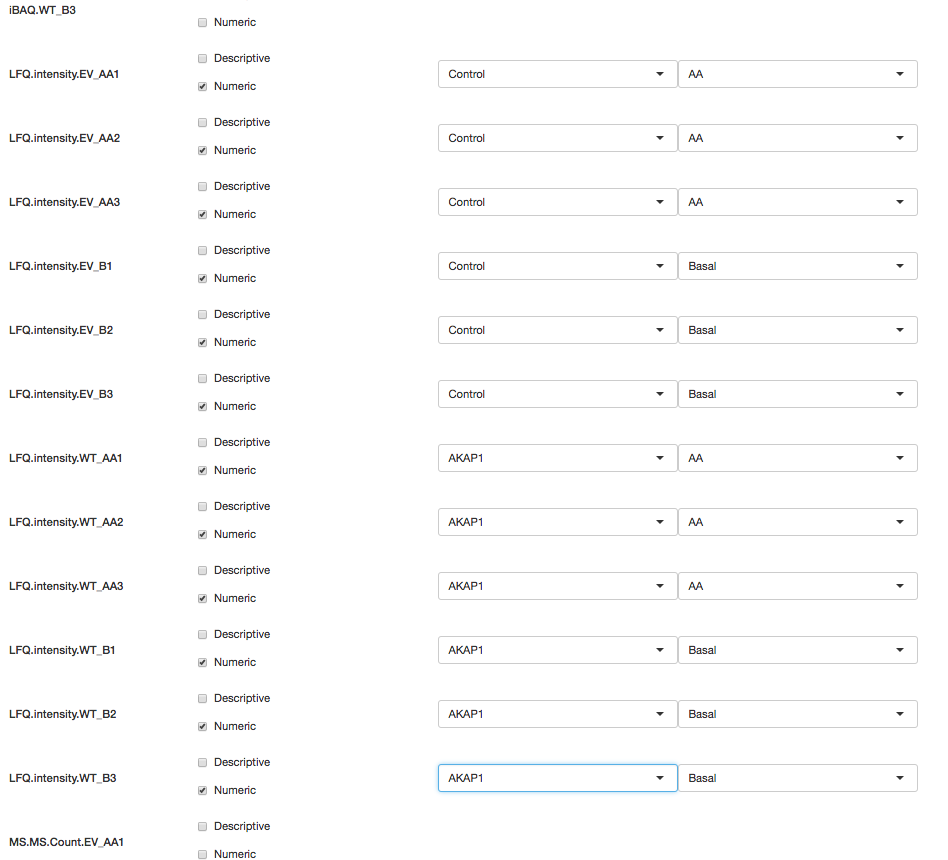

Fig. T2-E

Once all columns of interest have been annotated, we can apply the experimental design on the data set by clicking “Apply design and continue” button in the bottom right corner of the page (Fig. T2-F). This takes us to the next page for exploration, quality control and analysis.

Fig. T2-
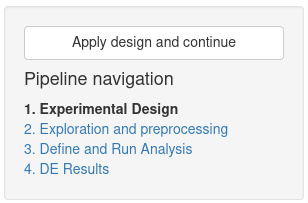
F

### Module: DATA EXPLORATION AND PRE-PROCESSING

In this module, we explore properties and trends in the dataset using a series of diagnostic plots. The grey toolbar at the left side of the screen contains tools for pre-processing the dataset including log-transforming, filtering, batch removal, imputation and normalisation of data. The main body of the screen, on the right, displays reactive diagnostic plots reflecting any such changes. The user may begin by log-transforming the data and plotting the expression levels of their favourite proteins of interest or positive controls as shown in Fig. T3-A and B, respectively.

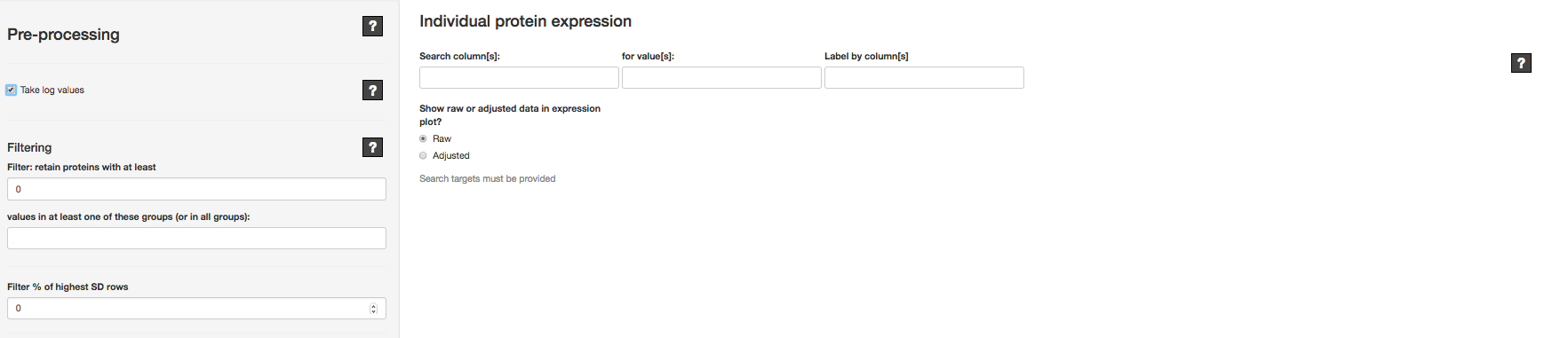

Fig. T3-A

In the main body of the screen, the “Individual protein expression” section allows the user to display expression levels for selected proteins. Previously when annotating the dataset, columns were marked as “descriptive” or “numeric”. If a column is marked as “descriptive” then this column becomes available to search: for example, if the “Gene name” column has been marked descriptive, this may be selected in the “Search column(s)” field. Proteins that were pulled down during the AP/MS experiment will be listed and searchable by their gene name under “for value(s)”. If the user wishes to label proteins differently, a different column can be selected in the “Label by column(s)” field.

Once one or more proteins have been selected, the log2(intensity) values are plotted on the y-axis with individual scatter points reflecting column values and barplots reflecting group means. This value is proportional to the number of peptides associated with the protein of interest (marked on the x axis). In Fig. T3-B we show the expression of 3 proteins across the 4 experimental groups in the AKAP1 dataset. Where there is no measured intensity reported for a particular group (missing values, represented as *NA*) as in the case of ESYT2, no bar is plotted.

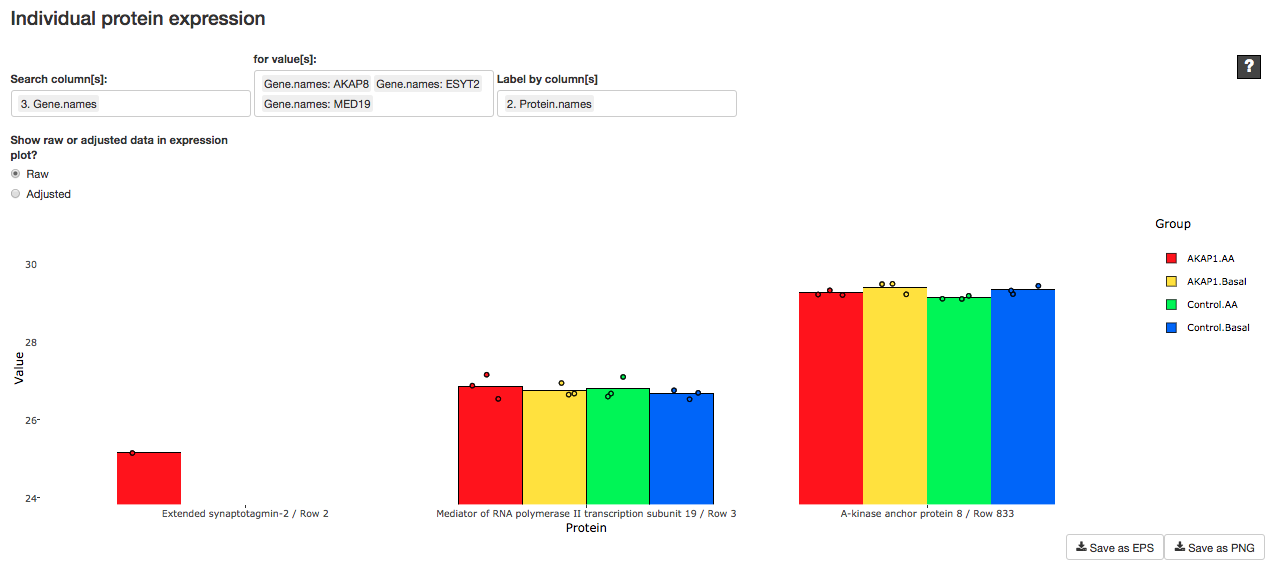

Fig. T3-B

#### Global diagnostic plots

The following diagnostic plots are generated on the fly on the log_2_ transformed dataset. Several of these plots are interactive, allowing users to hover over plotted objects to obtain more information about the samples or data points.

Sample distribution through boxplots (Fig. T4-B); and assess clustering and variability of groups through the clustering, variance (Fig. T4-C), and principal component analysis (Fig. T4-D) plots. The ranked heatmap of the data provides additional insight into sample distributions not available through the boxplot, as well as the relationship between intensity and missingness per sample (Fig. T4-E). The intensity versus missingness plot further allows the user to evaluate the structure of missing data by group, assisting in the selection of imputation method if necessary (Fig. T4-F). Finally singular value decomposition plots provide information as to sources of signal in the dataset, helping the user evaluate whether imputation should be done prior to- or post-normalization to minimize added noise (Fig. T4-G) (Karpievitch et al. 2012).

Users are able to determine the prevalence of missing values in their data through the “Missingness per sample” plot, displayed in Fig. T4-A. This graph allows the user to view what percentage of proteins are missing in each replicate trial.

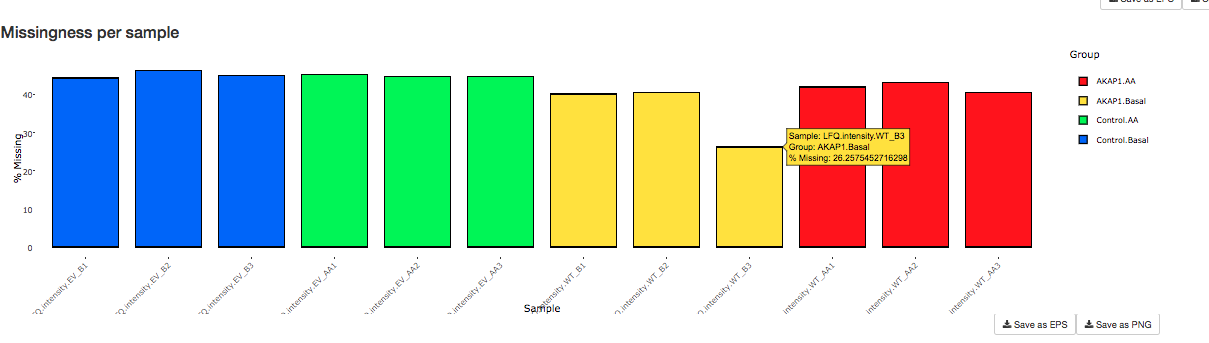

Fig. T4-A

Sample boxplots illustrate the distribution of intensity values for each sample, coloured by experimental group (Fig. T4-B). Under the assumption that differential signal exists only in a small number of proteins in the dataset, samples are expected to have similar medians; a departure from this suggests technical variation (as opposed to biological variation) and may require sample normalisation. Differences in missingness structure can also impact the appearance of distributions.

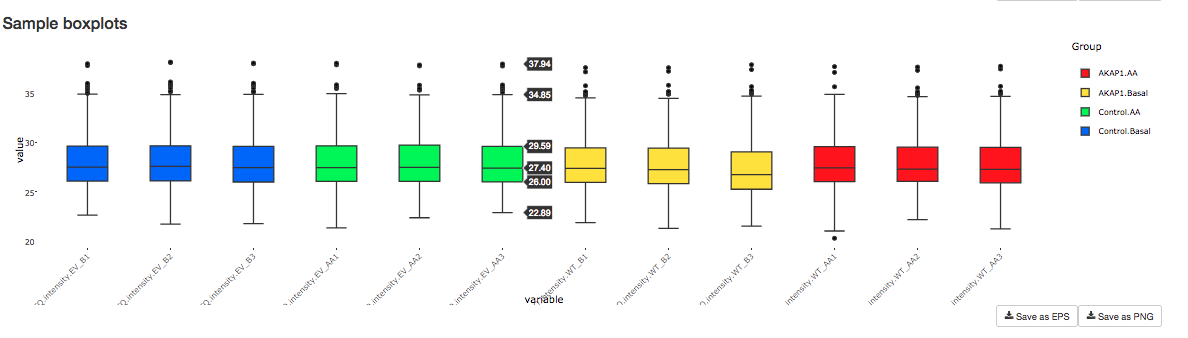

Fig. T4-B

Users may assess clustering and variability of groups through the hierarchical clustering and variance plots. (T4-C). As samples in the same group are expected to be biologically more similar than those in other groups, we hope to see samples clustered by group.

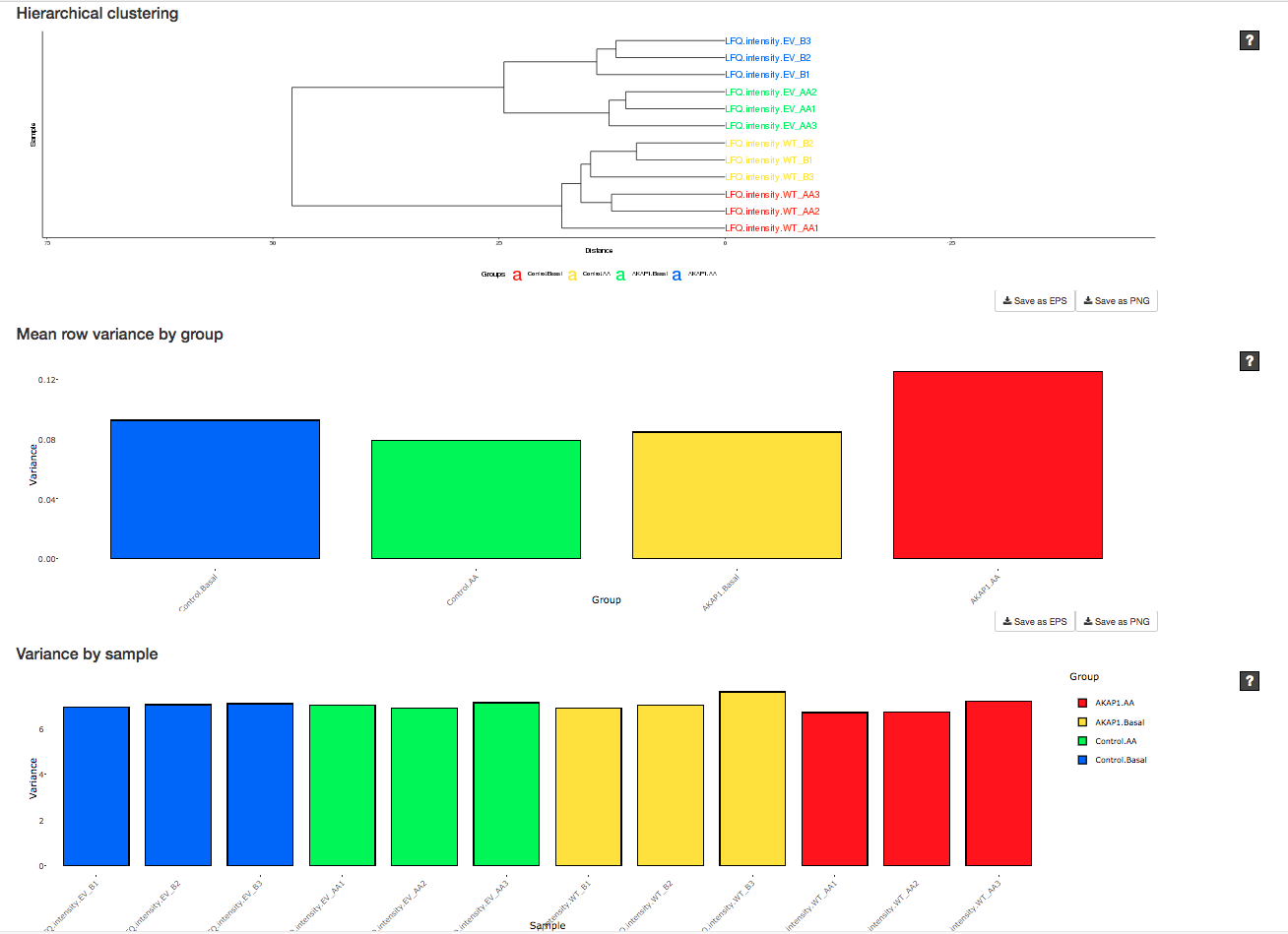

Fig. T4-C

The PCA plot (T4-D) displays a dimensionally-reduced representation of samples while maximising the amount of protein intensity signal retained. Similar to the hierarchical clustering plot in T4-C, replicates from the same experimental group should be close together in the PCA plot, while different groups should be separate from one another.

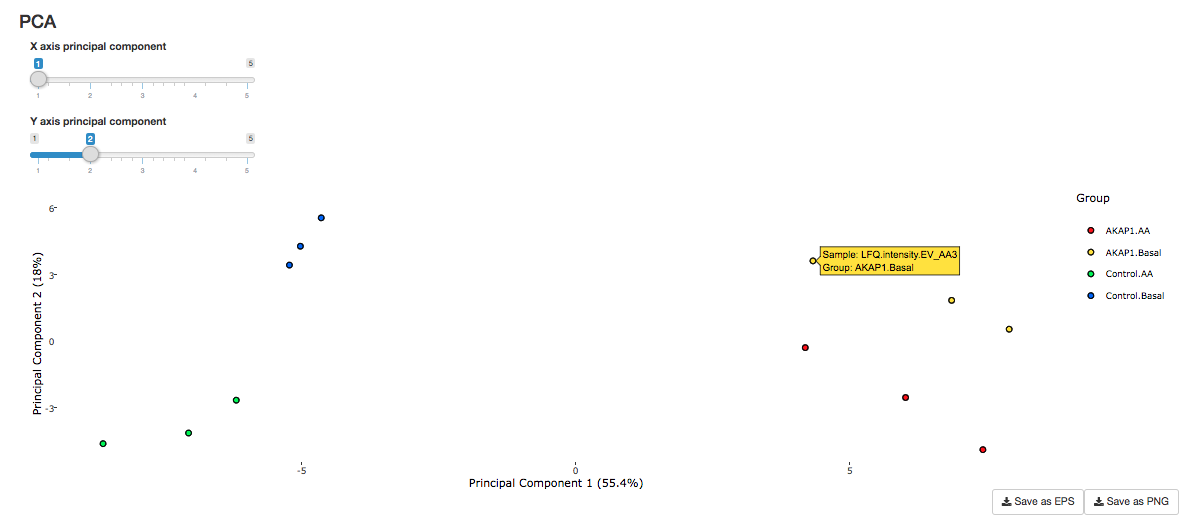

Fig. T4-D

An intensity rank heatmap of the data provides additional insight into sample distributions not available through the boxplot, as well as the relationship between intensity and missingness per sample (Fig. T4 E-F, top). Each value is coloured according to its global intensity rank, with rows ordered according to mean row intensity or number of missing values, as selected. This allows the user to observe the spread of the data without being confounded by missing values.

The “Intensity vs Missingness” (T4 E-F, bottom) plot further allows the user to evaluate the structure of missing data by group. This helps the user decide on an appropriate method of imputation, if necessary: Random Tail Imputation, for example, assumes that a positive relationship exists between a protein’s intensity and its probability of being detected, resulting in more missing values towards the lower end of the intensity range. This is what we will later use to process the example interactomics dataset.

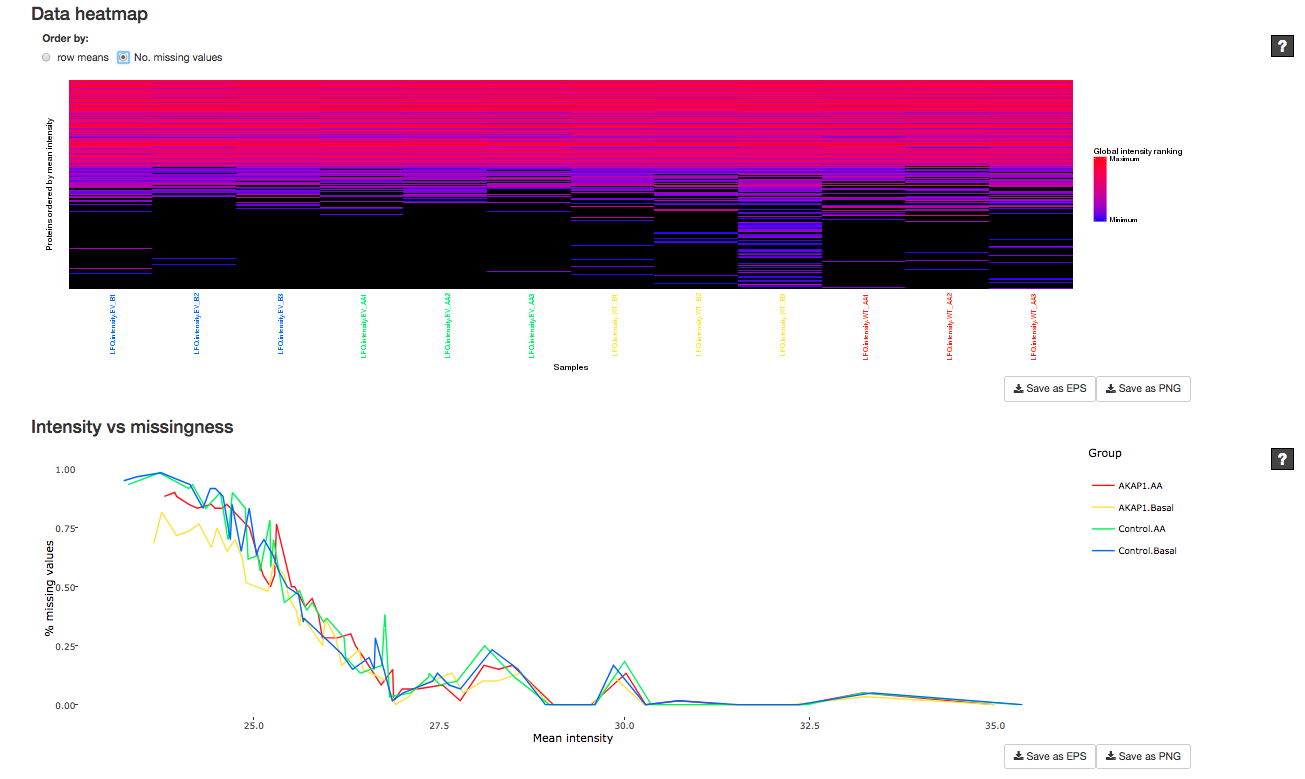

Fig. T4 E-F

Finally, singular value decomposition plots (Karpievitch et al. 2012)

separate sources of signal in the dataset, helping the user evaluate whether imputation should be done prior to- or post-normalization to minimize added noise (Fig. T4-G).

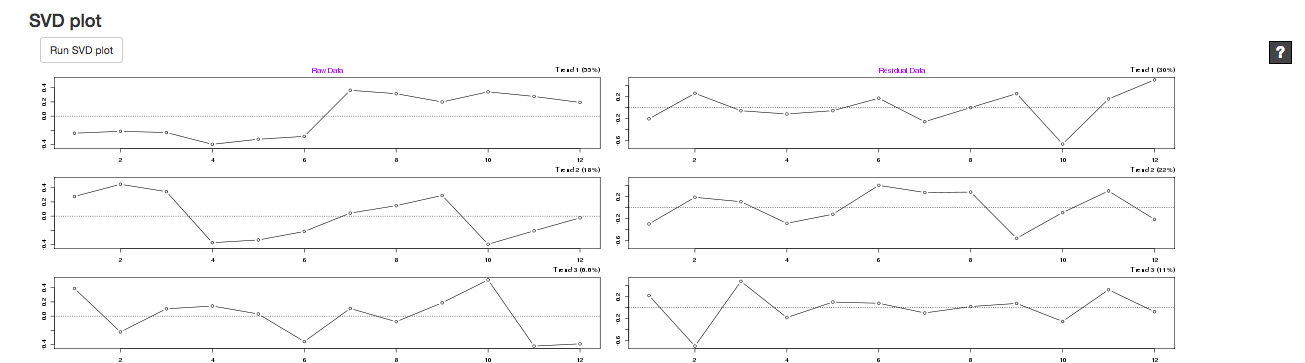

Fig. T4-G

In the SVD plot, ideally trend 1 in the raw data column (Fig. T4-G upper left) should show a clear separation in trend between replicates in the control groups and the replicates in the bait protein pull down groups. As the computation time for this plot is non-trivial, it does not update in response to changes in the pre-processing pipeline; the user must click the “Run SVD plot” button to refresh it.

#### Filtering to reduce missingness

Any pre-processing steps applied to the data are instantly reflected in diagnostic plots, helping the user understand the impact of these changes.

To put things into context, in an AP/MS experiment, each record (protein) in a replicate (sample) will either possess a positive log_2_ intensity value (if peptides of a protein are observed) or an *NA* value (if no peptides for a protein are observed/identified). A protein observed in one replicate may or may not be observed in other replicates. Proteins present in too few replicates may not be useful from an analysis perspective, and it may be more useful to remove them.

Thunderbolt allows us to do this using the fields under “Filter: retain proteins with at least ___ values in at least one of these groups ____”. The first field refers to the minimum number of replicates per group in which we require a protein to be observed. The second field allows us to specify the experimental groups to check. In experiments such as this, a protein is naturally often not observed under the empty vector control; in such a case, we may only wish to filter proteins absent groups where a bait protein is present.

In Fig. T5-A, we opt to retain only records observed in at least 2 of 3 replicates in the two experimental groups AKAP1.Basal (or WT_B) and AKAP1.AA (or WT_AA).

“Filter % highest SD rows” allows the user to remove proteins that are highly variable, as they may be more noisy. Removing this noise may improve our analysis, but is optional.

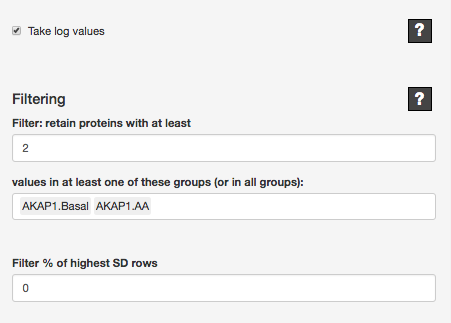

Fig. T5-A

The changes can be observed immediately in the “Missingness per sample” plot (Fig. T5-B), where the highest % missingness across samples has dropped from 40% in the unfiltered dataset (Fig. T4-A) to 15%. Similarly, all other plots are updated to reflect the change.

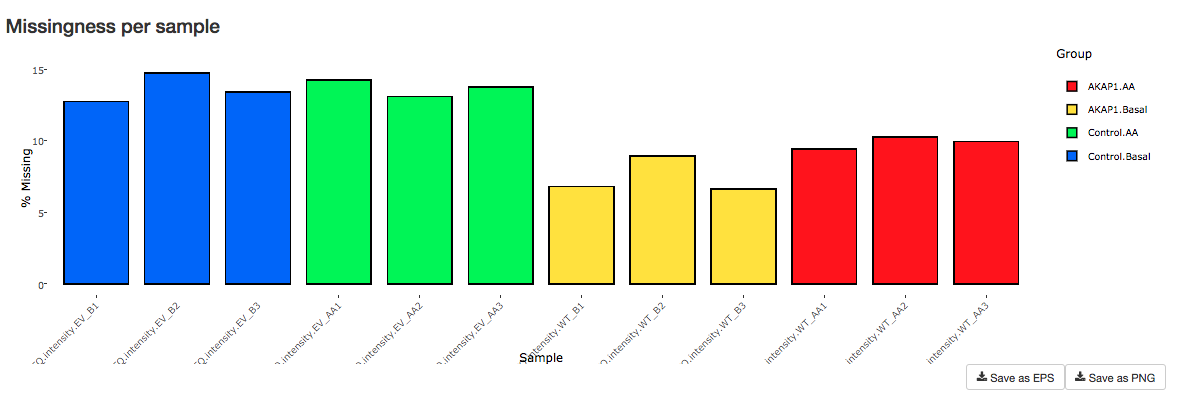

Fig. T5-B

#### Imputation and normalization

Next we may choose to impute and normalise the data using appropriate methods. The following fields are shown in in Fig. T6-A:

Normalisation options are provided both before and after imputation, including median normalisation, in which the median of each sample distribution is forced to the sample value, and quantile normalisation, in which corresponding quantiles of each sample are forced to the same value.

Next, multiple options are provided for imputation of missing values. Statistical analyses, including the differential expression analysis offered by Thunderbolt, must often operate on multiple values per group per test. Where a group contains 0 or 1 observations, such as the empty vector control in an IP experiment, it may be counter-productive to remove such proteins; indeed, ideally an interactor would be observed only when the bait protein is present. Hence, we may wish to impute missing values.

Thunderbolt currently permits two types of imputation: Random Tail Imputation (Deeb et al. 2012; Hubner et al. 2010) (appropriate where lower-intensity proteins are more likely to be missing), which draws random samples from normal distributed down-shifted and variance-reduced from the sample distribution; and k-Nearest Neighbours imputation (Hastie, T. et al. 1999). Either of these may be selected under “Imputation method (if any)” (Fig. T6-A). Here we choose Random Tail Imputation.

The next two input fields (“Impute all missing in groups _____” and “Impute rows in groups ____ with fewer than ____observations”) allow the user to select which groups to impute and optionally whether to limit imputation to proteins with fewer than a minimum number of replicates in those groups. As imputation may add noise to the data, in our example we opt to limit our imputation to cases with fewer than two observations in a group.

As we have selected Random Tail Imputation, we may choose whether to apply random to use multiple imputation (MI) (Rubin, 1987) in our DE analysis and how many imputations/analyses to perform in parallel (Fig. T6-A, “How many imputations”). As Random Tail Imputation involves random sampling, the significance of tests on imputed proteins may differ from run to run; MI allows us to perform imputation multiple times and combined DE analyses to produce a final set of more robust statistics. We may also choose to alter the parameters (Fig. T6-A, “Shift” and “Width”) (Deeb et al. 2012) of the distribution from which our imputations are sampled, which is reflected in a reactive histogram.

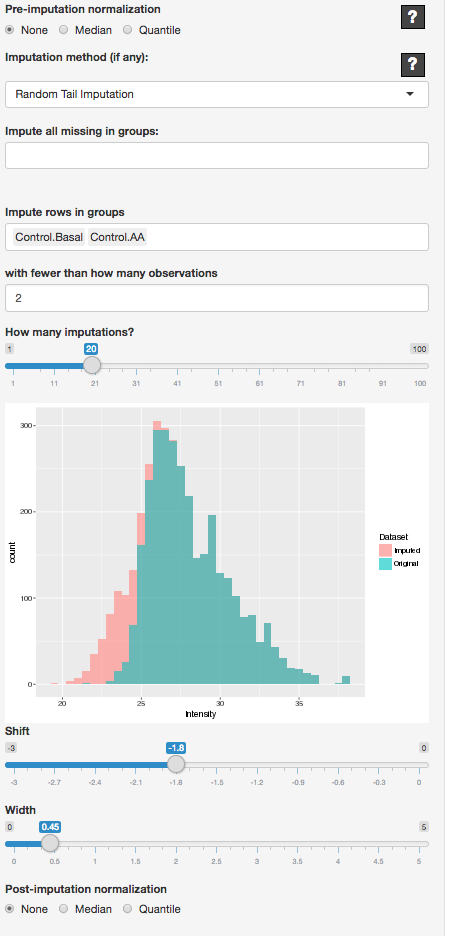

Fig. T6-A

The resulting imputed data are reflected through the diagnostic plots. The following plots (Fig. T6 B-H) are shown as examples. Note that if multiple imputation is used, only one set of imputations is used to generate the plots.

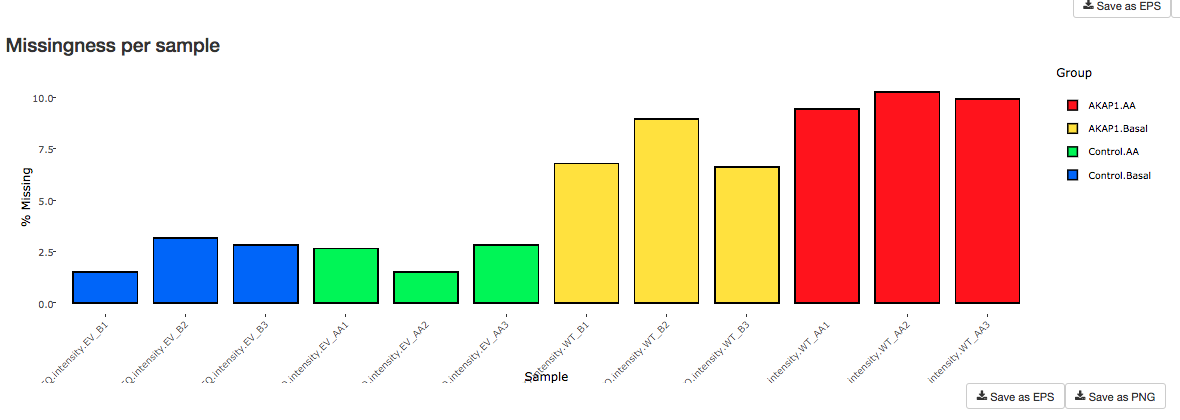

Fig. T6-B

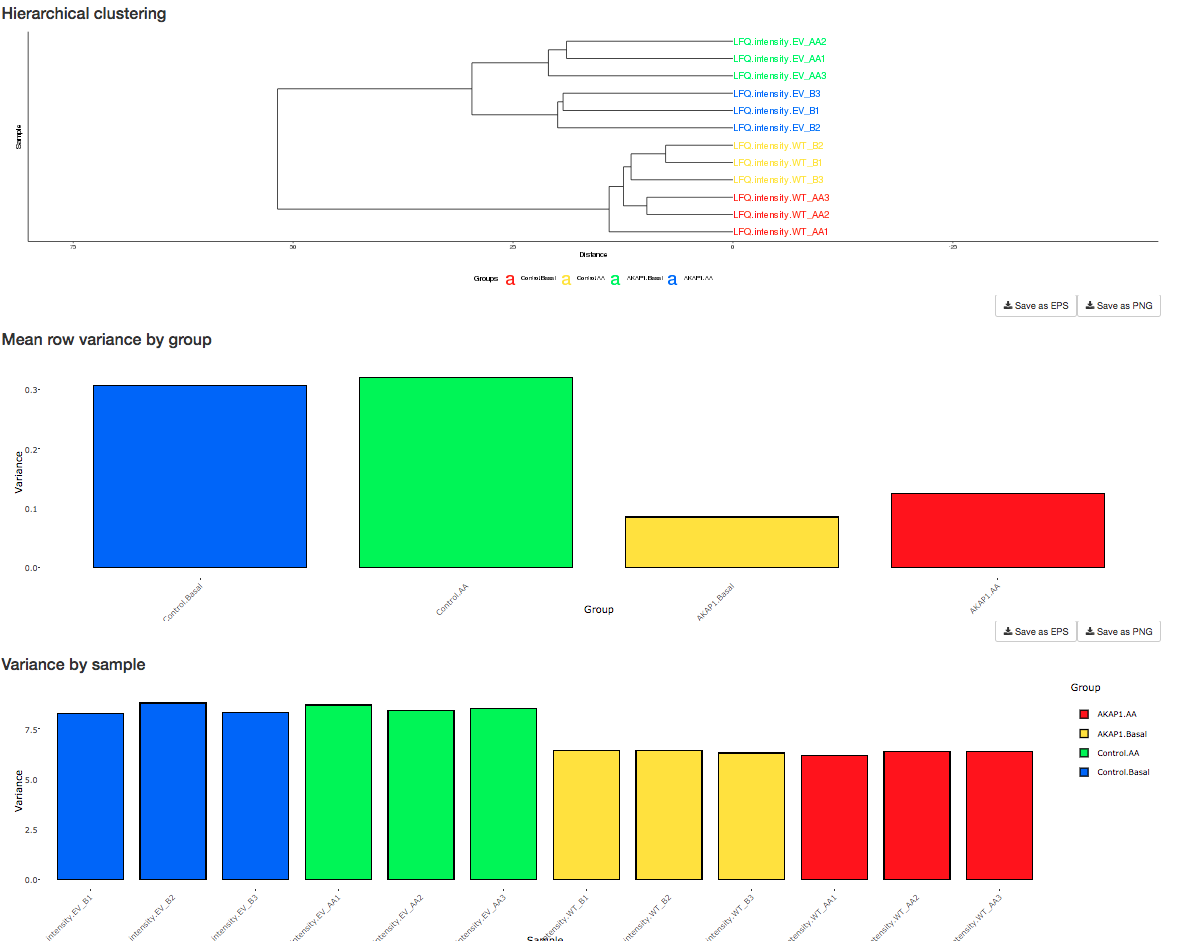

Fig. T6 C-E

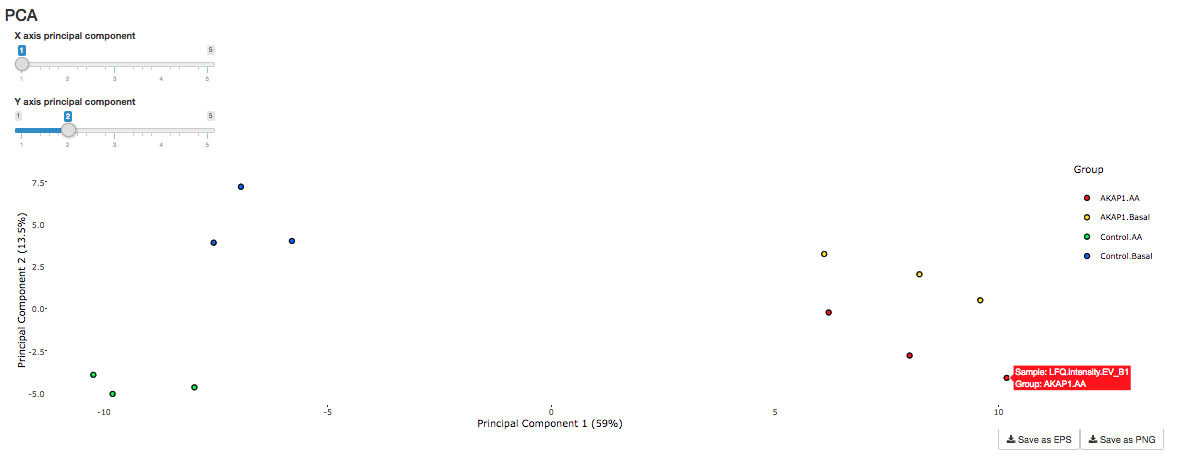

Fig. T6-F

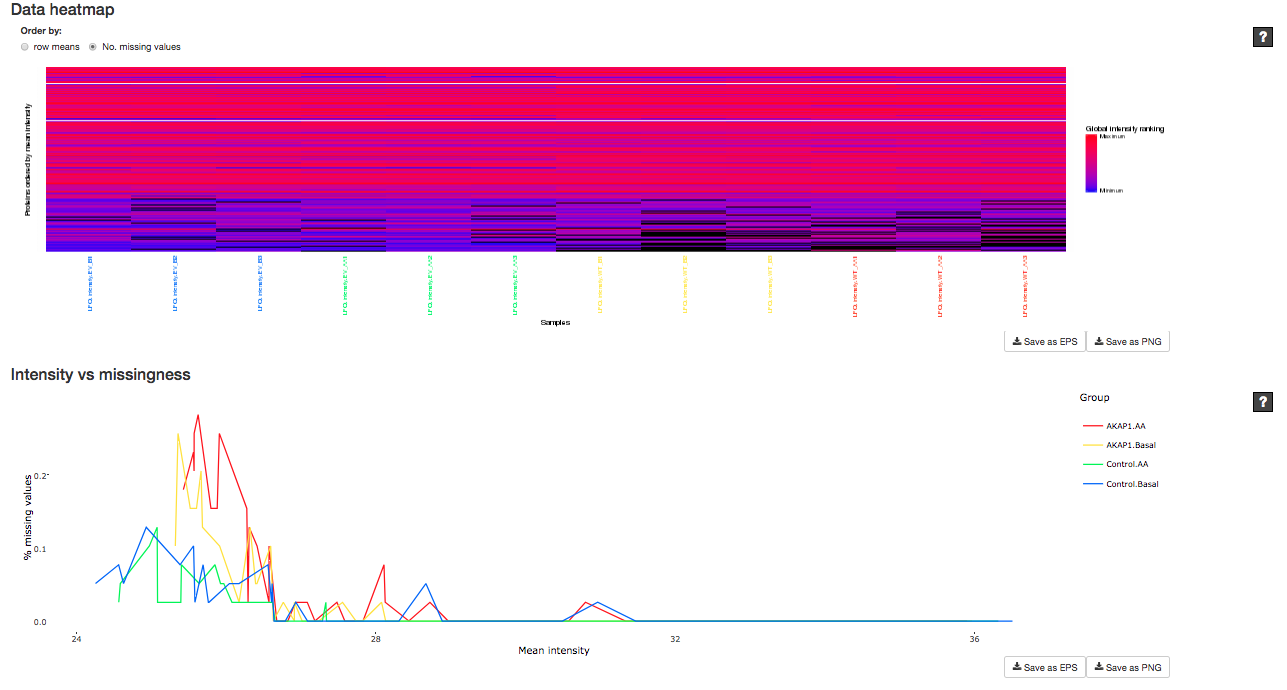

Fig. T6 G-H

### Module: DE ANALYSIS

Once the user is satisfied with their dataset, differential expression (DE) analysis can be carried out. In this module, we first specify which groups we wish to compare using the moderated t-test as implemented in limma (Ritchie et al. 2015). As shown in Fig. T7-A, we can enter these comparisons under the “Current contrasts” section, first entering a name and then a contrast formula (typically in the format “Group_A – Group_B”, but more complex contrasts are also supported). Once satisfied, we can “Analyse” the data.
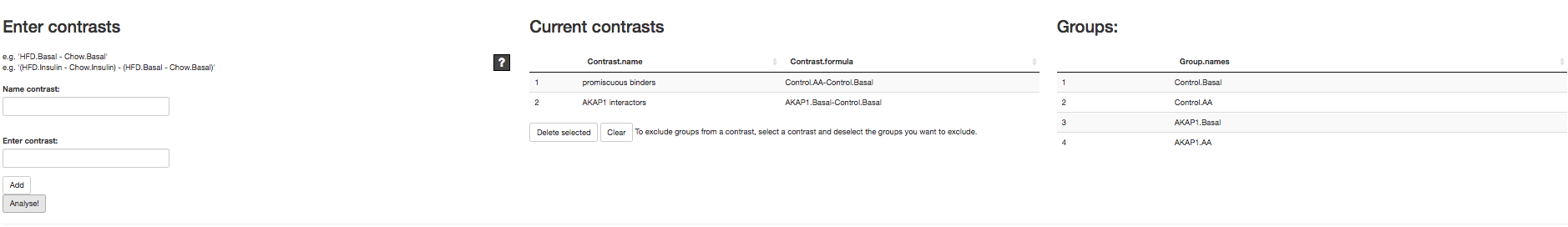

Fig. T7-A

In our example, we perform two comparisons. First, we compare the two control groups to obtain promiscuous binders (ideally, there should be nothing altered between these conditions; if something is significantly altered, it is a false positive which needs to be eliminated from the results).

Under the second comparison, labelled “AKAP1 interactors” we compare the bait pull down to empty vector under basal conditions; this should provide us with AKAP1 binders.

Fig. T7-B shows the DE results section. In order to visualise the DE analysis results, four global plots are presented; each of these may be saved locally in either the .png or .eps file format. The “T-Values” plot shows the distribution of t-statistics from the moderated t-test. The plots labelled “P-Values” and “P-Values (zoomed)” visualise the relative frequency of p values obtained from the DE analysis. Under the null hypothesis (no truly significant proteins), this is expected to exhibit a uniform distribution; thus, assuming some proteins are differentially expressed (or true interactors, in our IP example) we expect to see a largely uniform distribution with a spike of p-values close to 0. Finally, the “Volcano plot” visualises the significant proteins/genes/sites based both on the p-value and log-fold change significance thresholds specified on the left side of the screen.

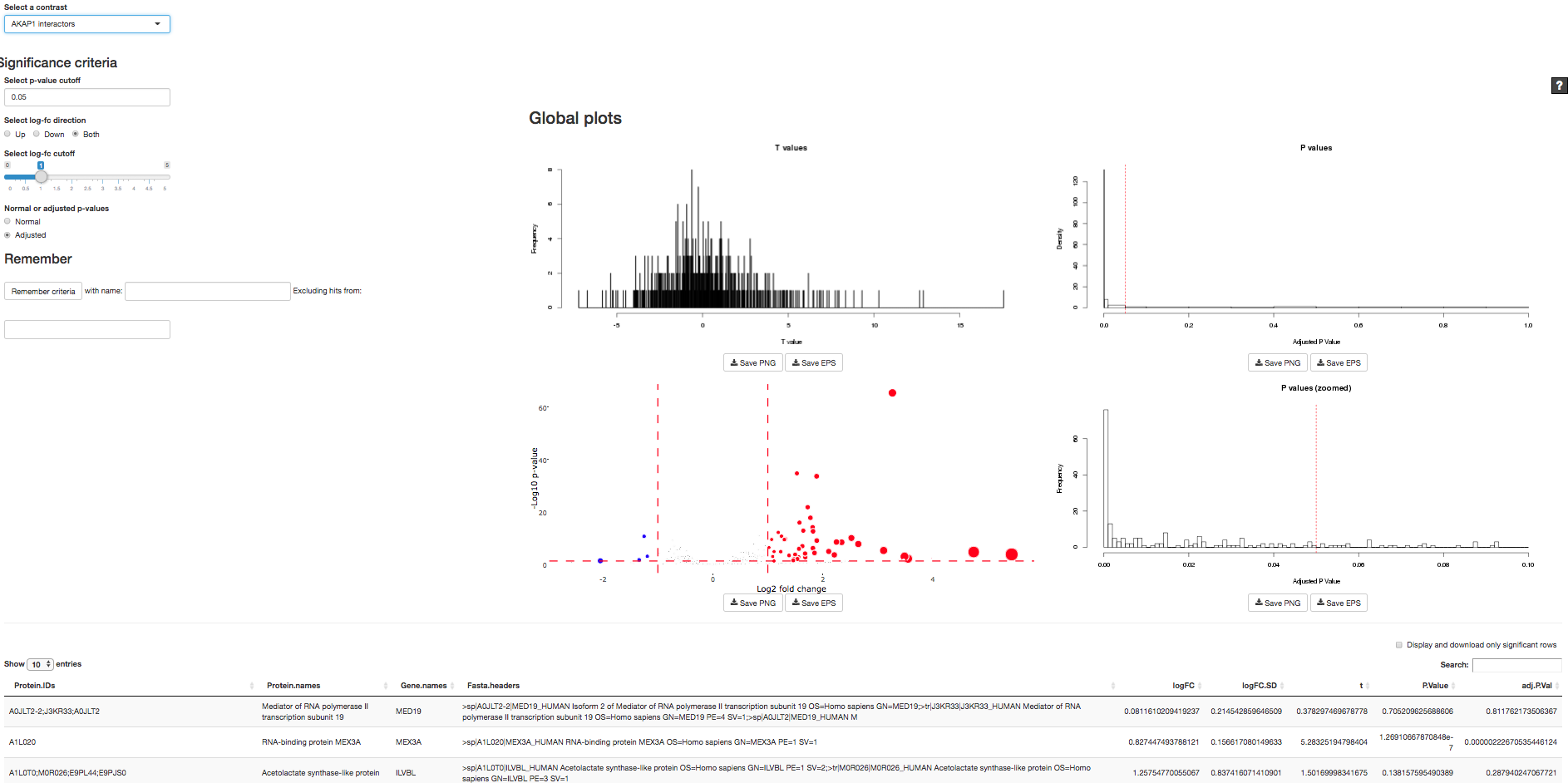

Fig. T7-B

The volcano plot can be inspected by hovering over the data points for additional details, as demonstrated in Fig. T7-C.

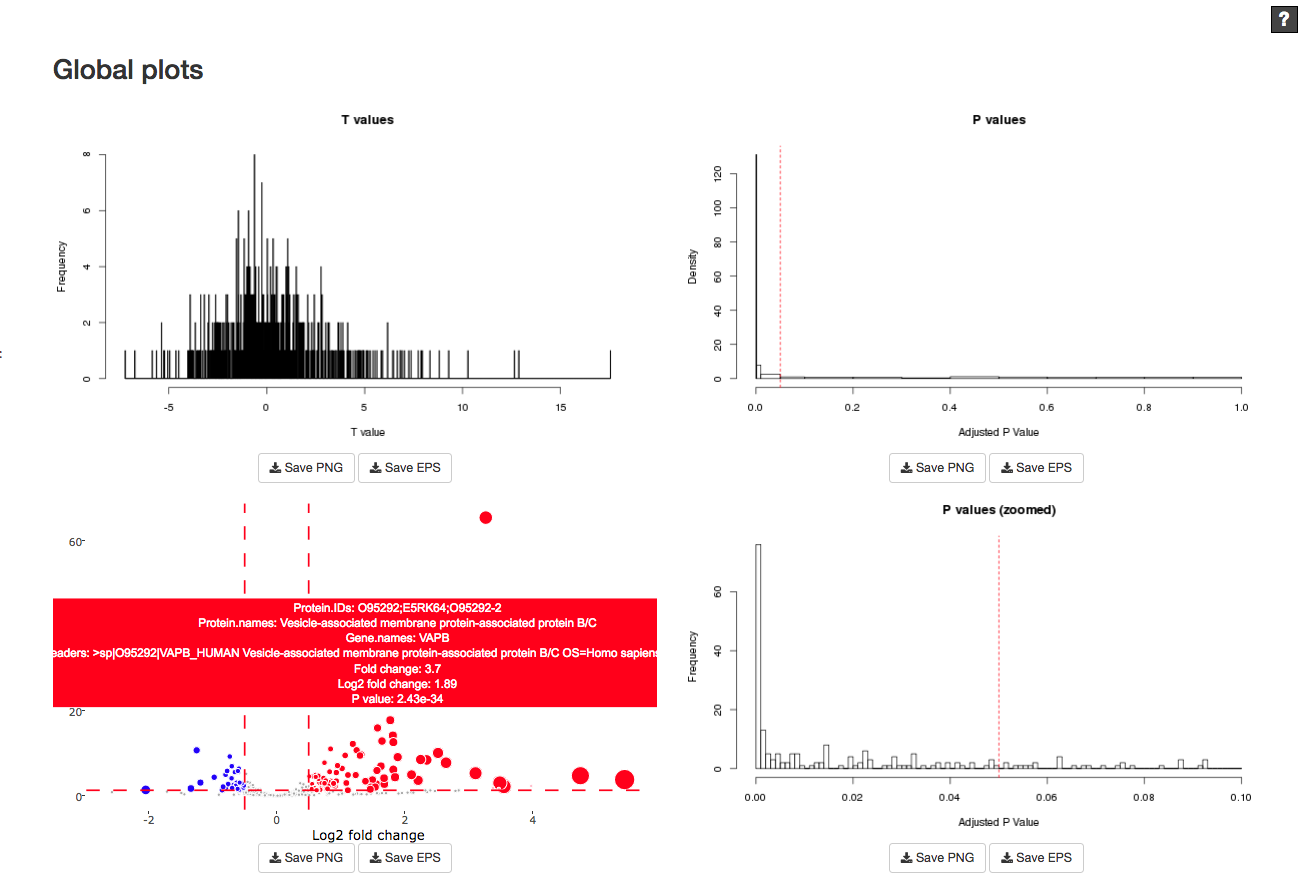

Fig. T7-C

The criteria by which a protein is determined “significant” can be adjusted using the sliders on the left of the screen (expanded in Fig. T7-D). Differentially expressed genes for a given comparison can be saved by “selecting a contrast” (Fig. T7-D), then specifying a “name” for the significant genes and pressing “Remember criteria” button.

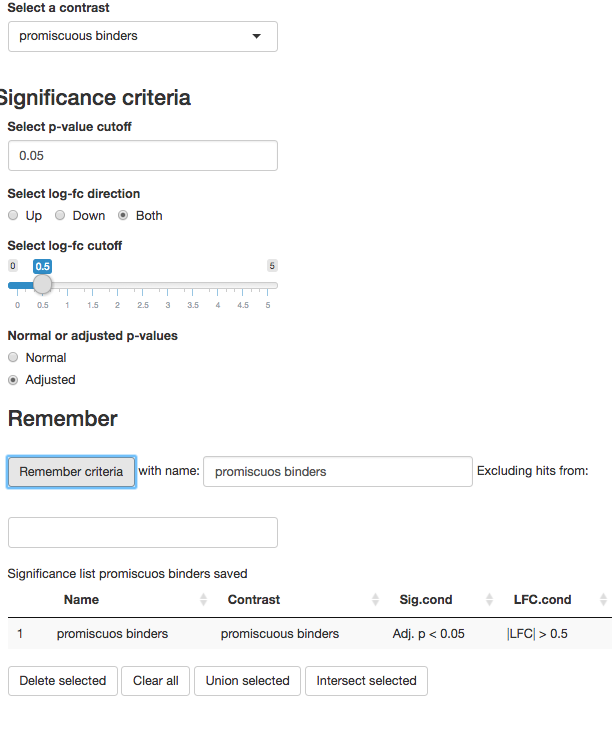

Fig. T7-D

Advanced options are available for generating lists of significant proteins by combining information from multiple comparisons. For example, a third list can be created as an intersection or union, or by excluding the contents of one list from another. In our example (Fig. T7-E), the promiscuous binder list is excluded from the AKAP1 interactome list to weed out the obvious false positives.

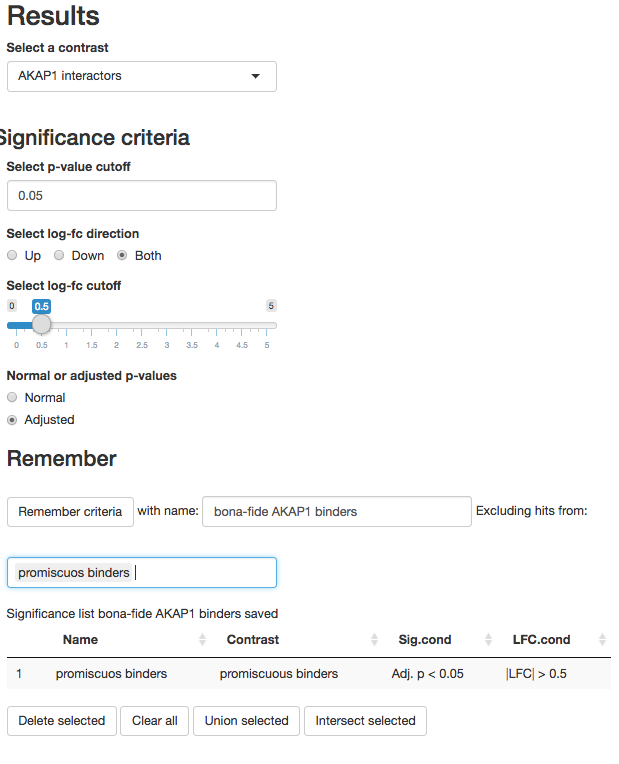

Fig. T7-E

The resulting proteins are displayed as a table, shown in Fig. T7-F. Optionally, only the significant genes can be displayed by checking the box “Display and download significant rows”. The table can be downloaded in a variety of formats.

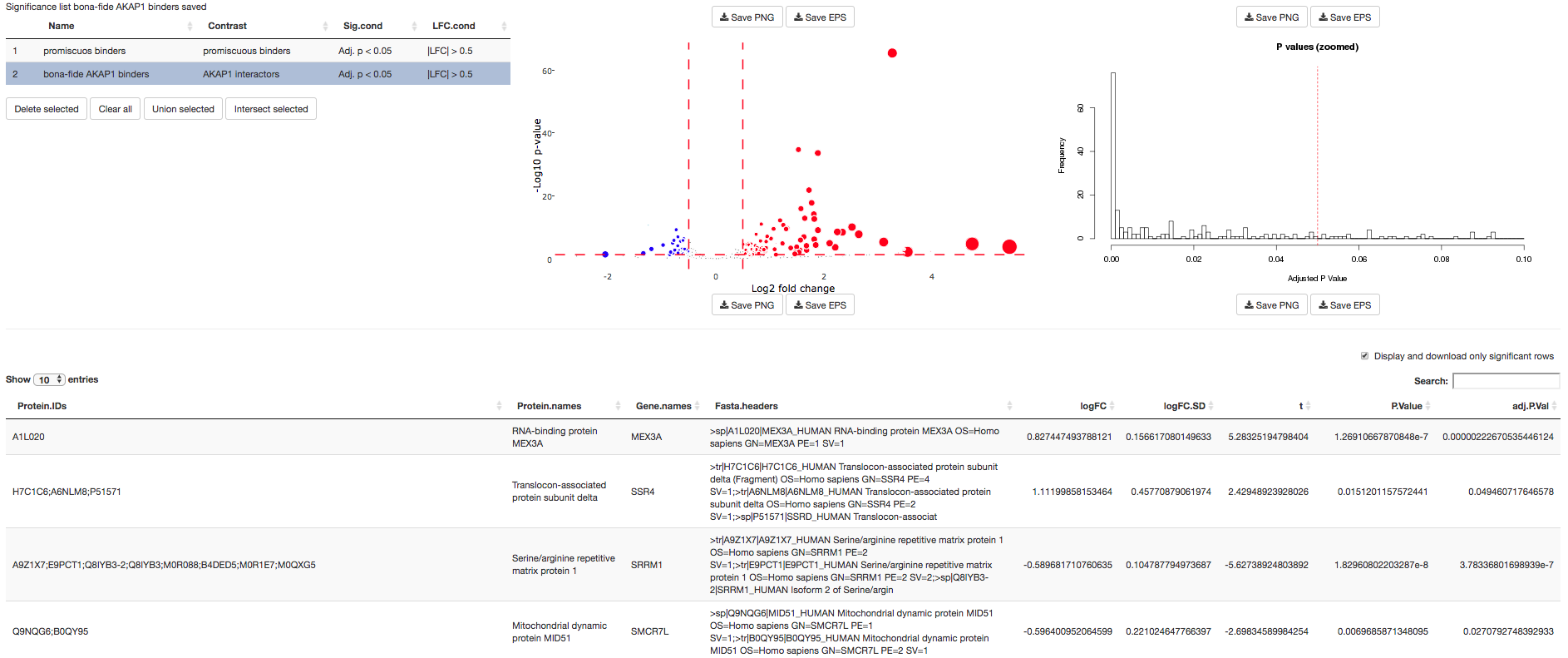

Fig. T7-F

### Module: FUNCTIONAL ENRICHMENT ANALYSIS (PATHWAYS)

The functional enrichment analysis module accepts a list of proteins and finds biological pathways whose proteins are statistically over-represented, or functionally enriched, in the list. The module tests each pathway listed in the KEGG PATHWAY database (Kanehisa & Goto, 2000) using Fisher’s exact test, and can operate on uploaded lists or those generated by the DE module.

We click “Add new protein set” to begin. As we generated a list of proteins of interest in the previous section covering DE analysis, we select the second option “DE significance list” to retrieve it (Fig. T8-A). If the user instead has an existing list of proteins they wish to analyse, these can simply be uploaded in a file containing gene symbols, Uniprot IDs or Entrez IDs.

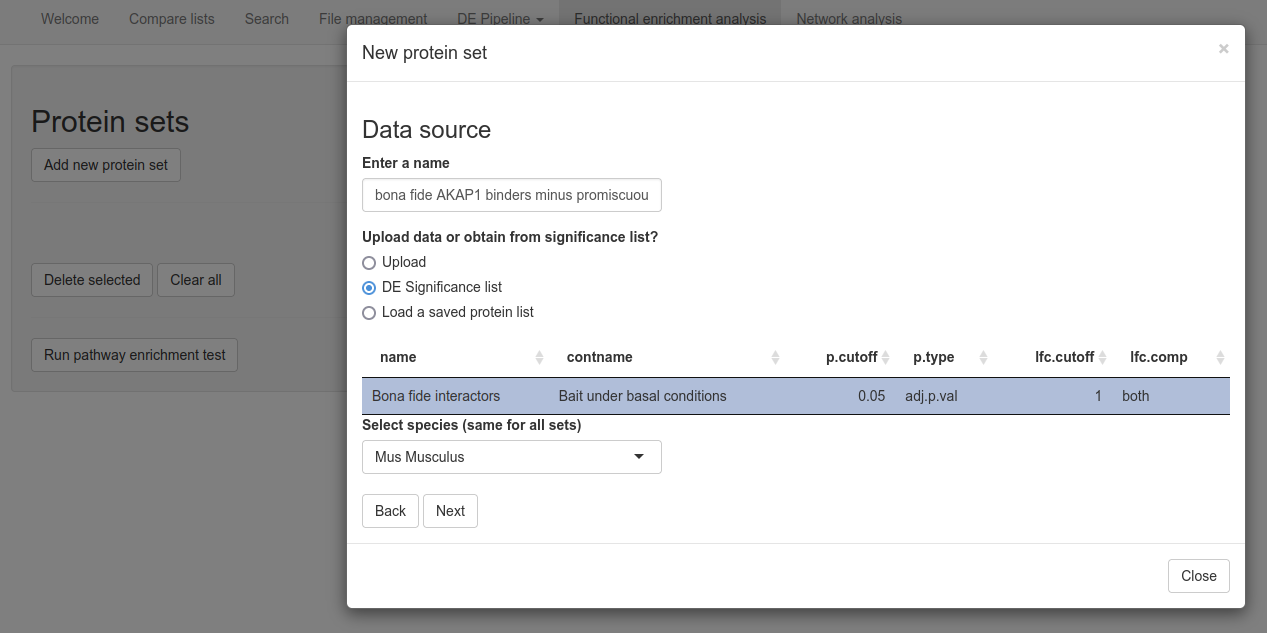
Fig. T8-B

After selecting the appropriate species, we click “Next”. On the next page, we are asked to select the appropriate column. In our example, we select the Gene names column (Fig. T8-B).

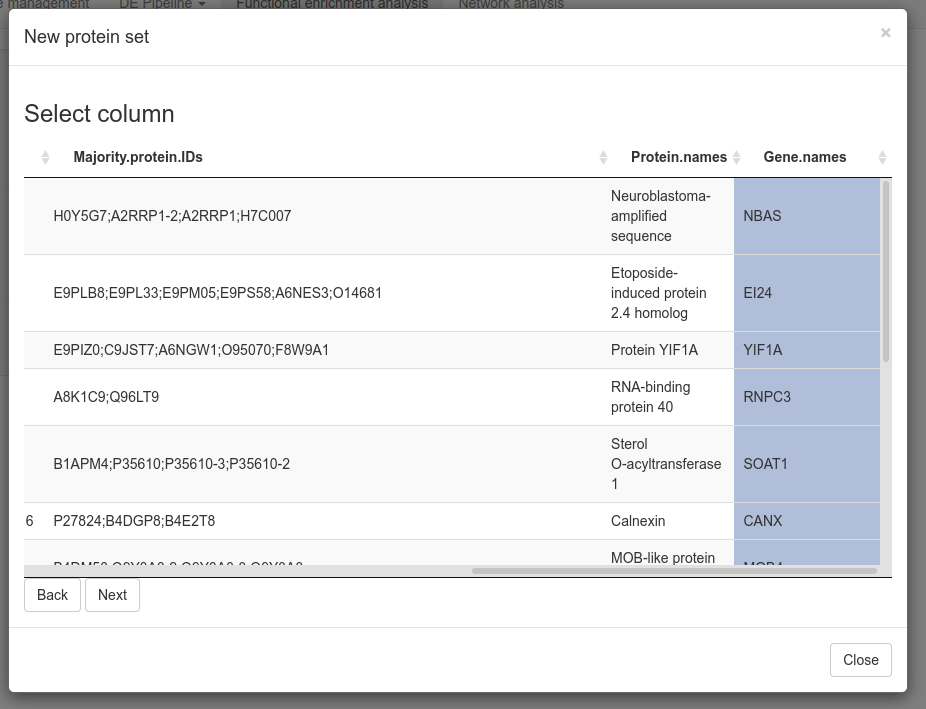

Fig. T8-B

On the next page, the user may specify the identifier type; the gene symbols in our data are converted to Entrez IDs internally as demonstrated in Fig. T8-C; this is required for compatibility with the KEGG database. If Entrez IDs are not displayed, a mistake may have been made on one of the previous screens. We click “Next” to return to the module’s main screen.

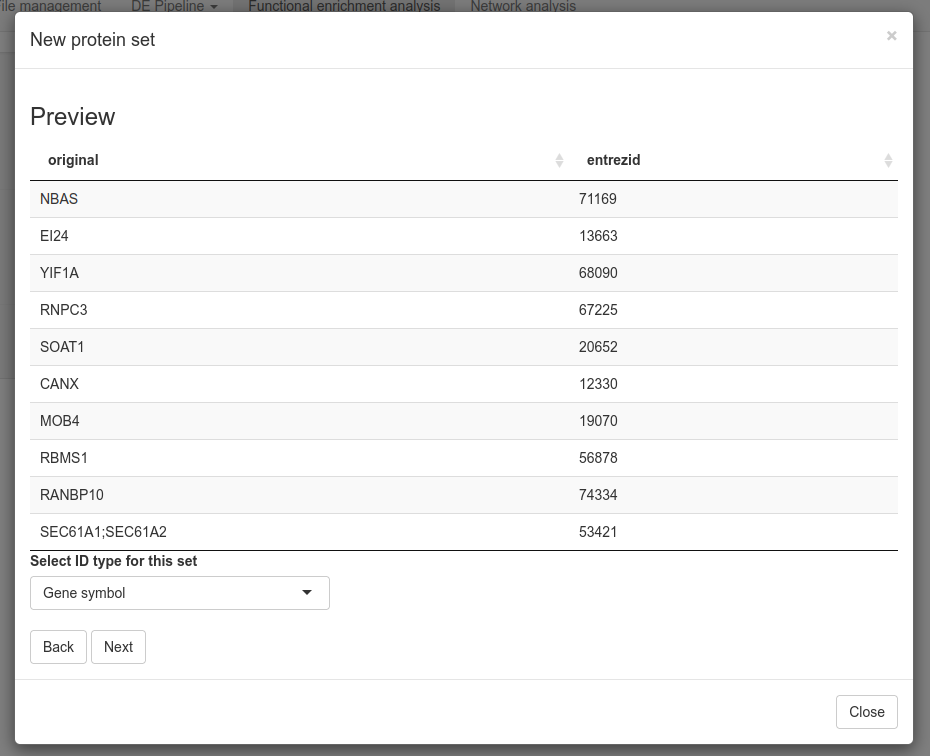
Fig. T8-C

Finally, the list of proteins can be tested for pathway enrichment by clicking “Run pathway enrichment test”. Results are available as a table or barplot, as displayed in Fig. T8-D. These may be downloaded in a variety of formats.

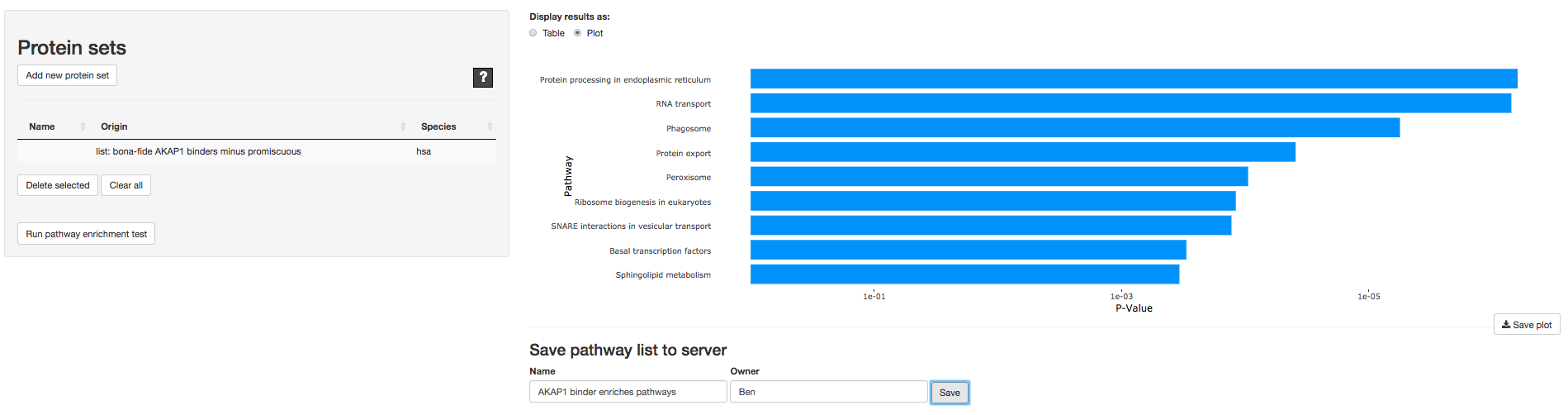

Fig. T8-D

The list of significant pathways may be saved to the server for use in the “Compare” module.

### Module: NETWORK ANALYSIS (STRING DATABASE)

The Network Analysis module allows users to visualise a list of proteins as a network graph, including known connections from the STRING database of protein-protein interactions (Snel et al., 2000). As in the functional enrichment module, the user begins by uploading or selecting a list of proteins and selecting a column of identifiers. The user must additionally specify a bait protein (if there is no specific bait, one can leave this field empty). In our case, the bait is “AKAP1” (Fig. T9-A) and prey proteins are taken from the significance list as gene symbols. If the user has uploaded a list containing a column specifying multiple bait proteins, this can be selected.

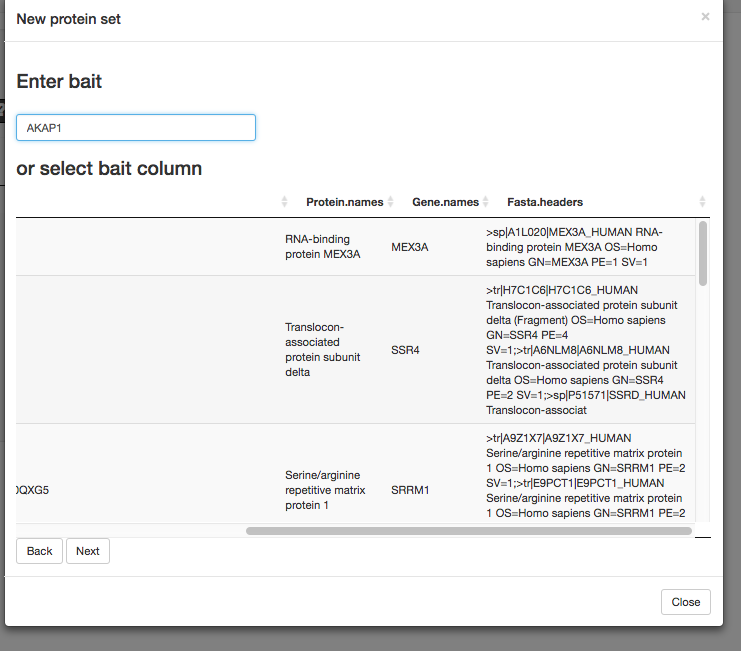

Fig. T9-A

After clicking “Next” to return to the main page of the module, we click “Create graph” to generate a simple graph showing AKAP1 in the middle connected to all DE proteins (Fig. T9-B). All visualisation is carried out using D3.js (Bostock et al., 2011).

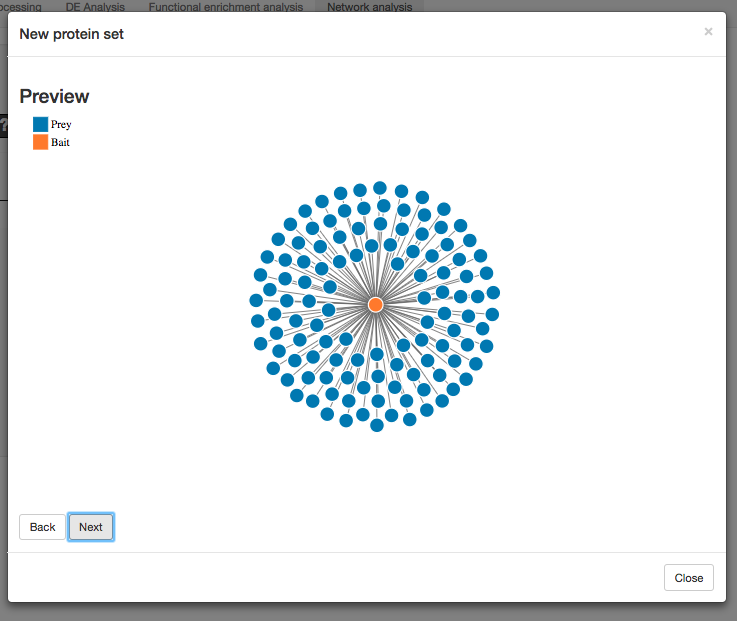

Fig. T9-B

By including prior protein-protein interaction knowledge from the STRING database, we can create more interesting networks such as those shown in Fig. T9-C and D. T9-C reflects known direct interactions between prey proteins, whereas D shows includes any proteins interacting with at least 2 prey proteins (in and outside of the given dataset).

Finally, basic network statistics such as hub and betweenness centrality metrics can be calculated and downloaded as shown in Fig. T9-E.

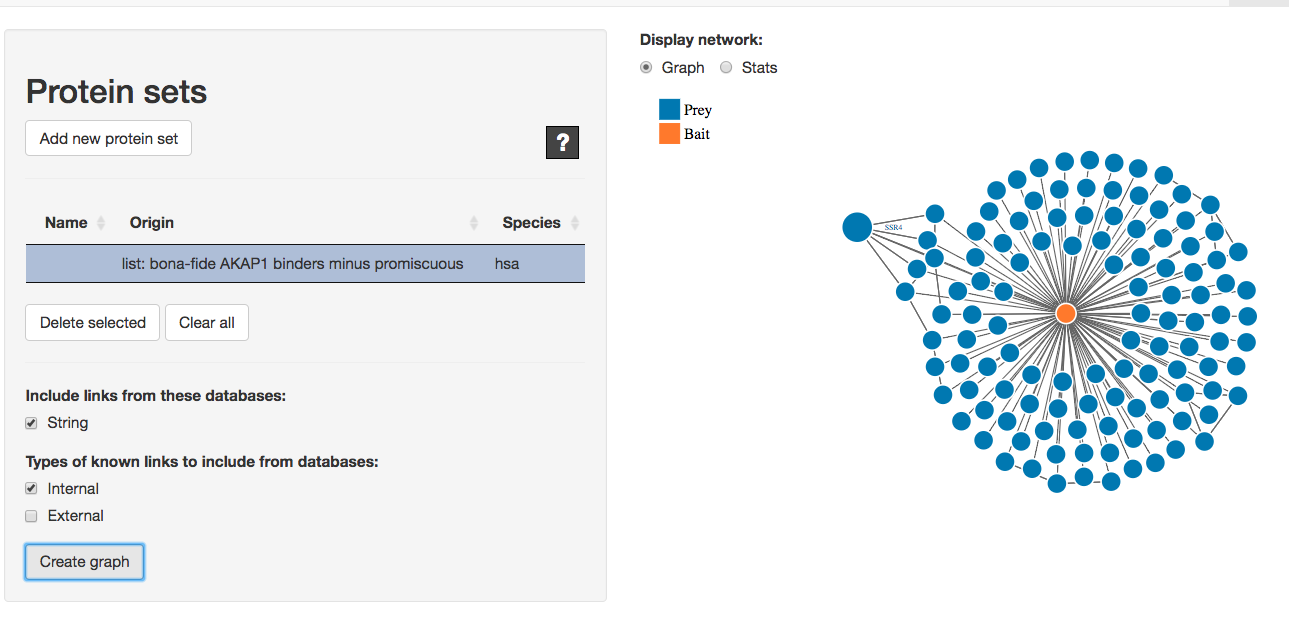

Fig. T9-C

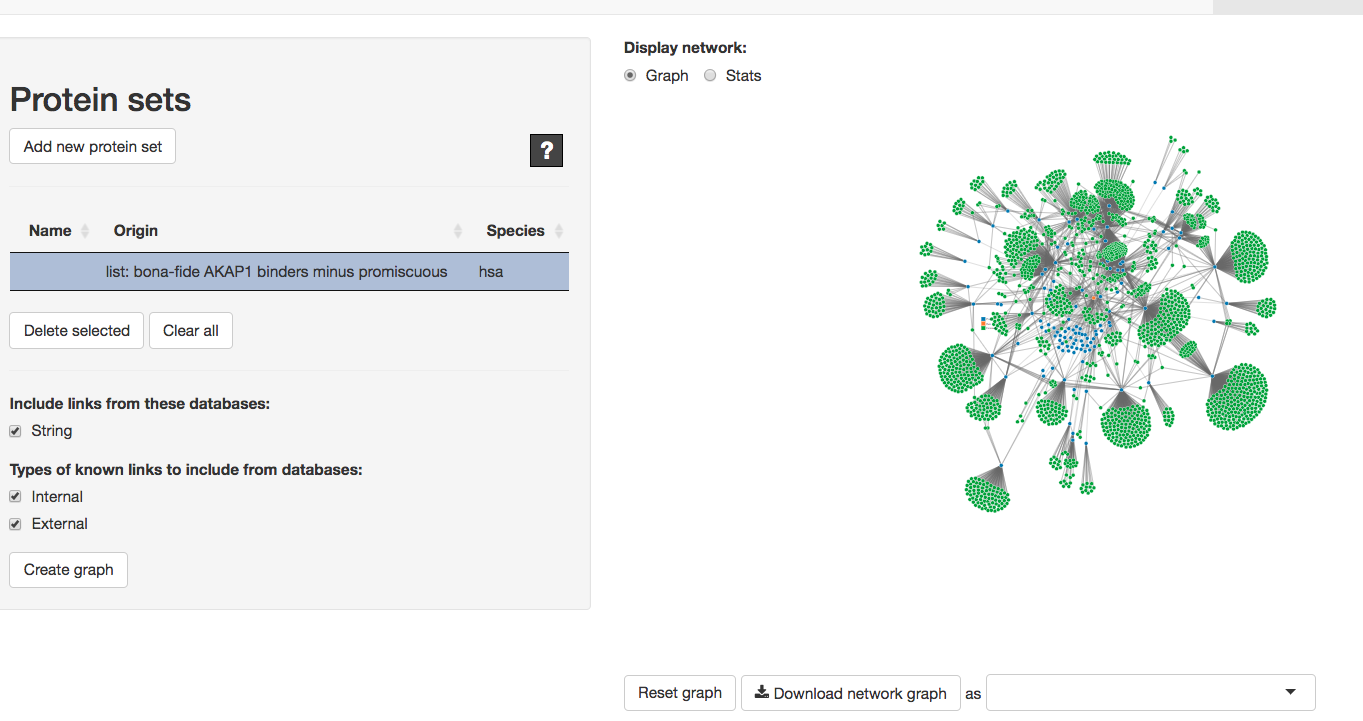

Fig. T9-D

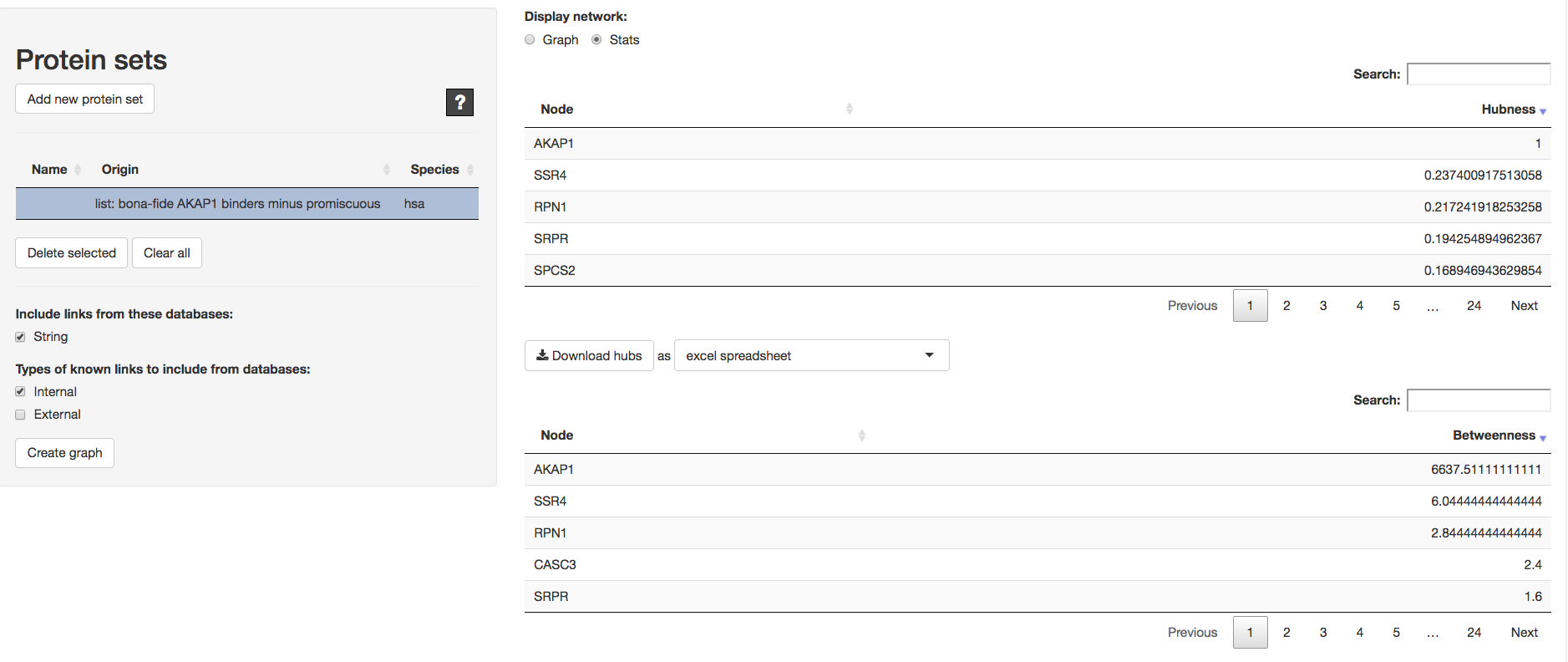

Fig. T9-E

### Module: COMPARE

This module allows users to compare multiple datasets (proteins, pathways or any descriptive information) in the form of lists.

Using proteins as an example, Fig. T10-A shows a Venn diagram of overlaps between three test lists of differentially expressed proteins previously saved into Thunderbolt. The output can also be displayed as an interactive heatmap (showing overlap counts or Jaccard indices) (Fig. T10-B) or as a table (Fig. T10-C) which can be sorted for records common across all datasets by toggling the column “in # lists. This table can also be downloaded in several formats including excel spreadsheet, csv, tsv and RData file.

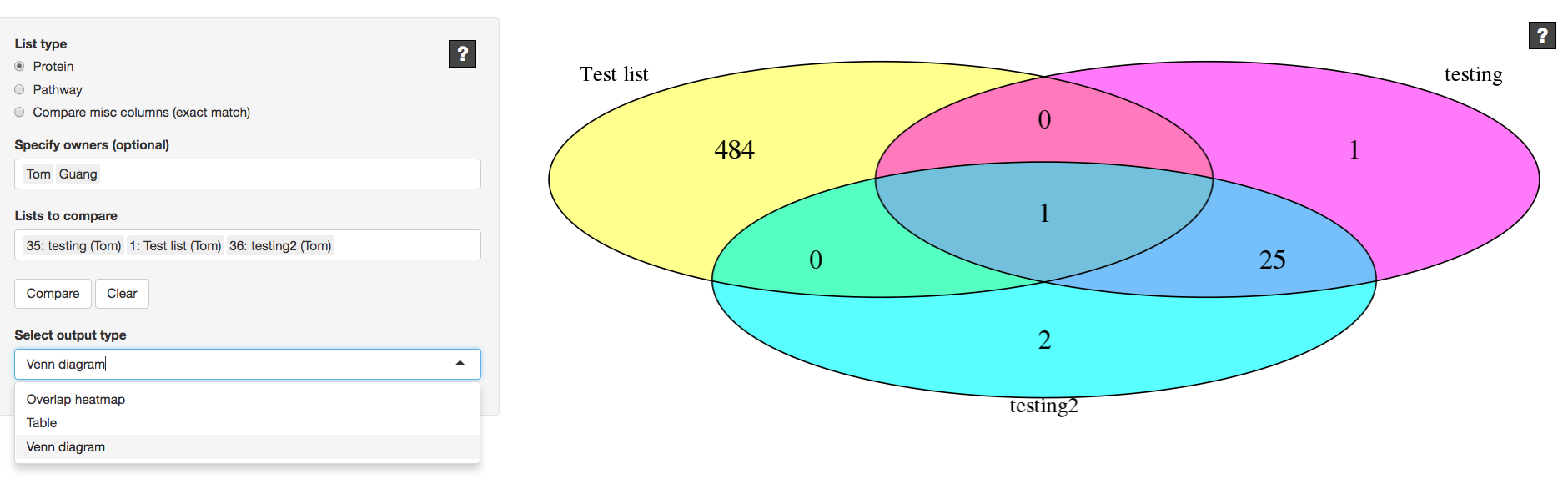

Fig. T10-A

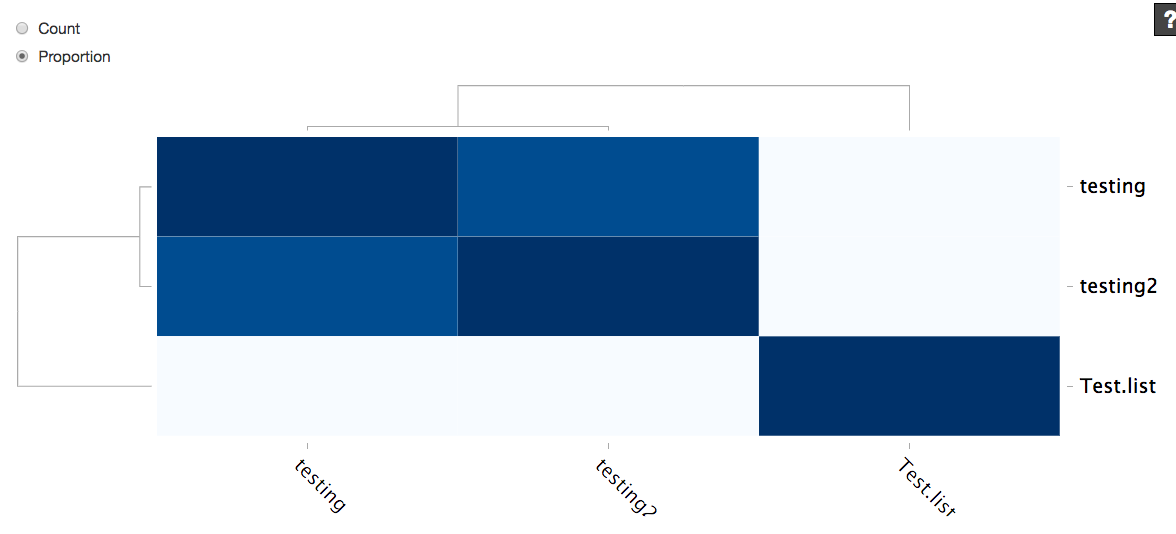

Fig. T10-B

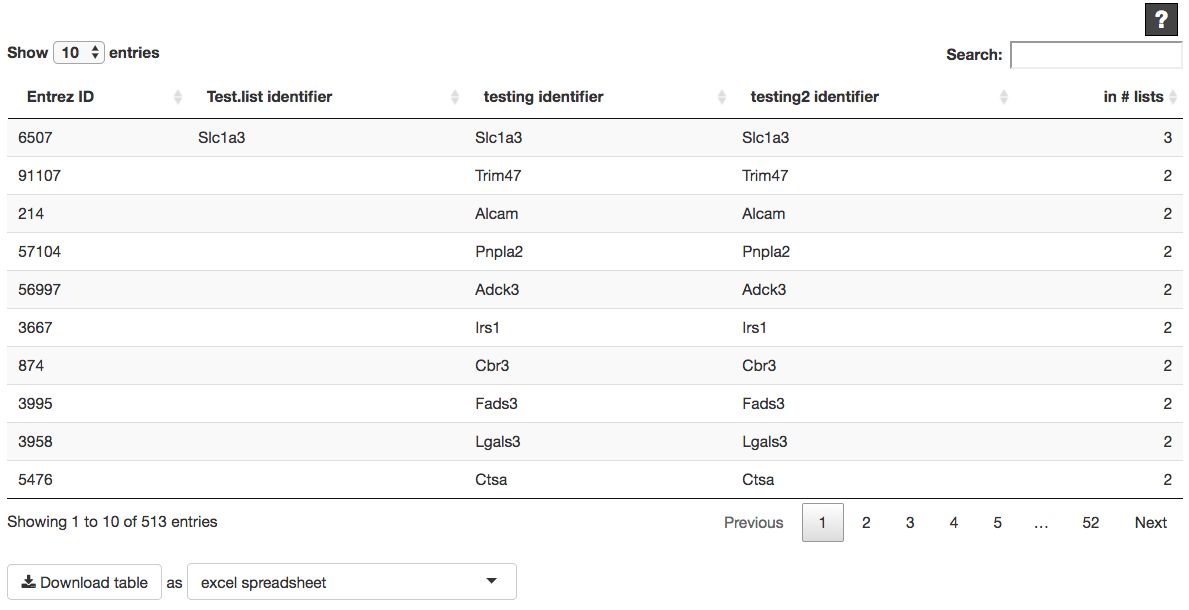

Fig. T10-C

### Module: SEARCH

This module allows users to search and collate results of selected proteins/genes of interest across multiple datasets. For example, if the user wants to know if their favourite protein *x* is DE in any of the proteomics or transcriptomics datasets to prioritize target validations, the user can search for *x* in any of the analysed and/or saved datasets in the Thunderbolt environment in this module. The output includes the associated statistics for *x* across all selected datasets, allowing the user to judge its importance from the displayed results.

For a given search type (differential expression analysis results or pre-saved protein lists), genes or proteins can be searched using gene symbol, Uniprot ID, Entrez ID or any other descriptive data column across multiple datasets. Users can also search for multiple proteins at once, or none: they can choose to filter their search solely through the third panel “Search by analysis output” without searching for any specific protein(s).

If the user chooses to search for analysed DE datasets (“Analysis results”), the third panel “Search by analysis output” allows us to filter our search based on log_2_ fold change and significance criteria (P-value or adjusted P Value) (Fig. T11-A). In this example, we search for the gene Akt1 in 3 datasets, as selected in the “Name” field under “Search by metadata”. Fig. T11-B lists datasets in which the gene “Akt1” can be found (2 out of the 3 selected datasets). Selecting one or both of these datasets, individual rows containing Akt1 are displayed in the table shown in Fig. T11-C. Associated statistics for Akt1 phosphorylation sites in the two datasets are displayed here. This table can be downloaded or saved as a protein list.

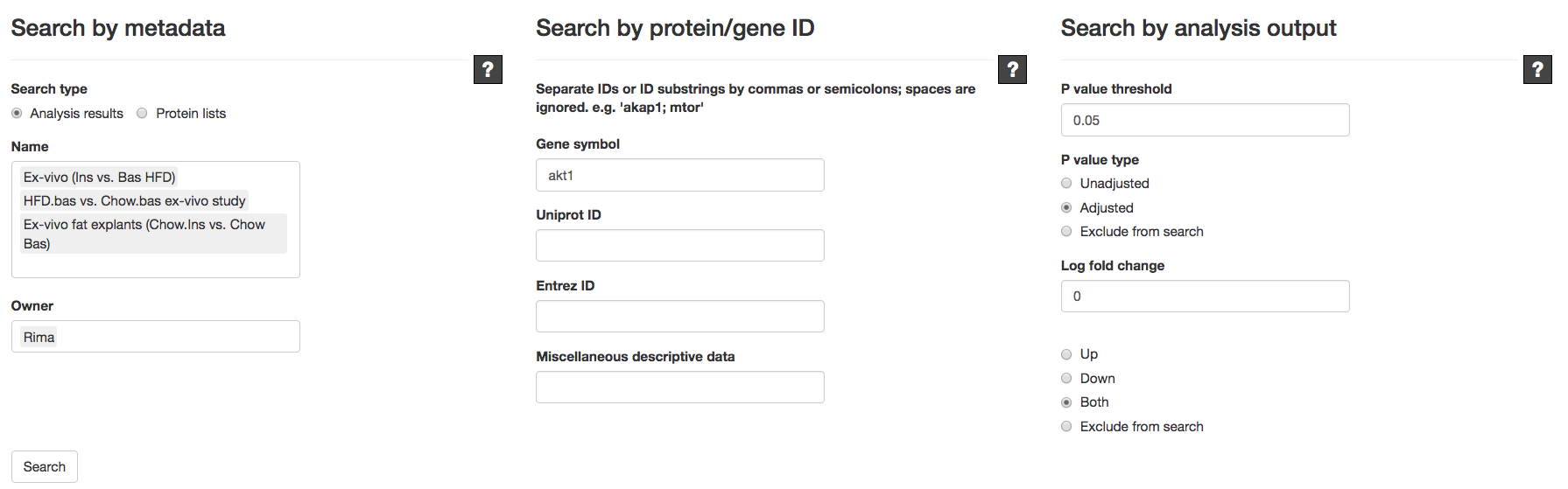

Fig. T11-A

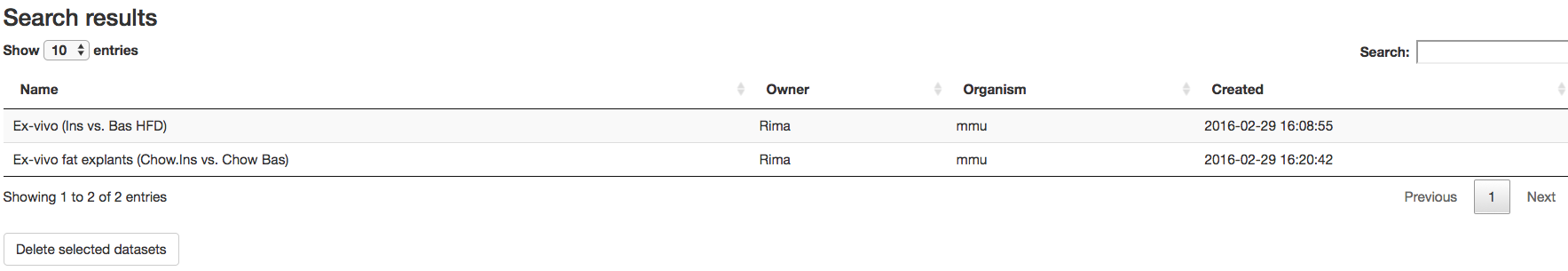

Fig. T11-B

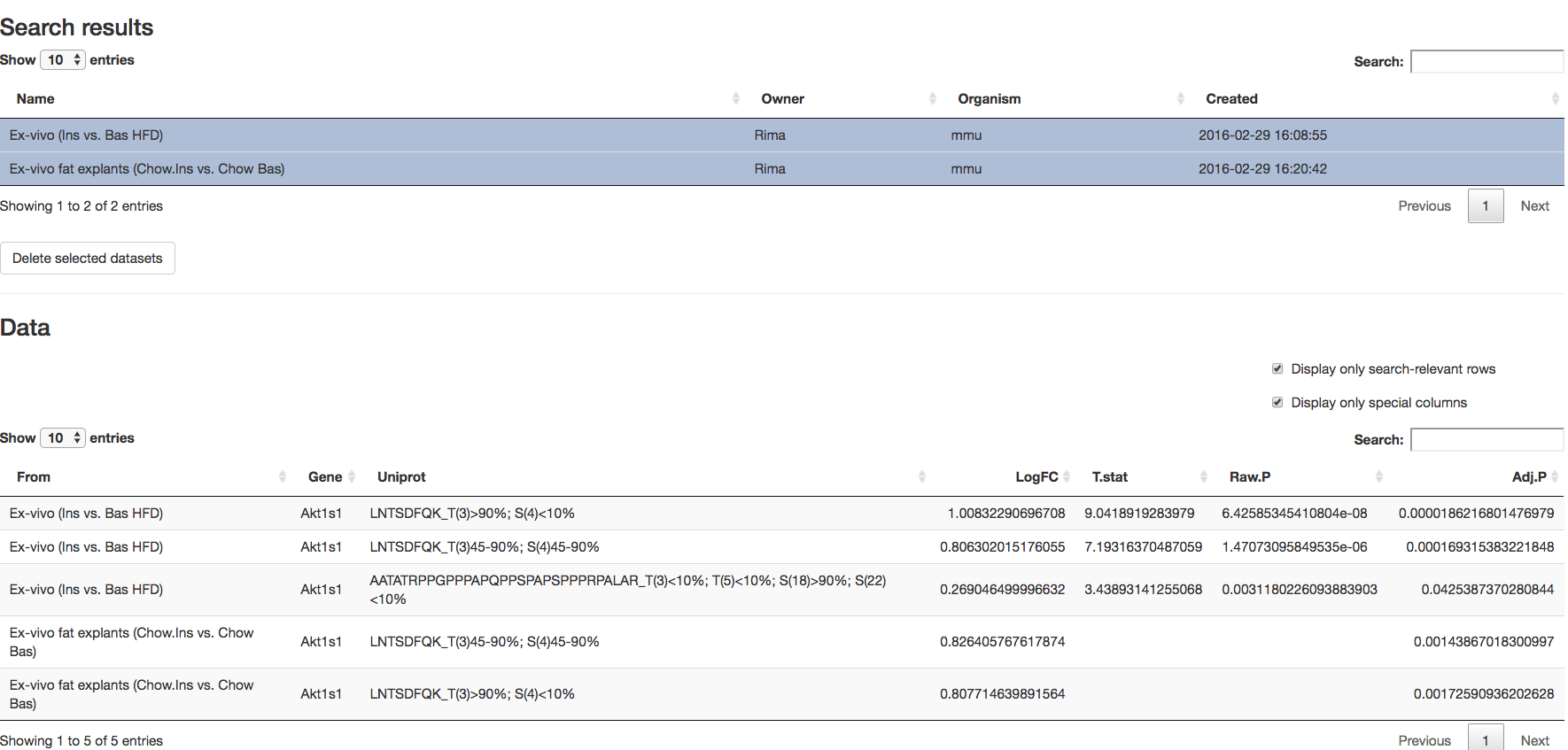

Fig. T11-C

If we wish to find all DE genes in the 2 datasets, we can simply leave the second panel “Search by protein/gene ID” empty and instead filter for DE genes using the third panel. Search results can be saved to the server as a protein list (Fig. T11-D) that can later be compared to other protein lists in the COMPARE module to estimate overlap of DE genes across experiments.

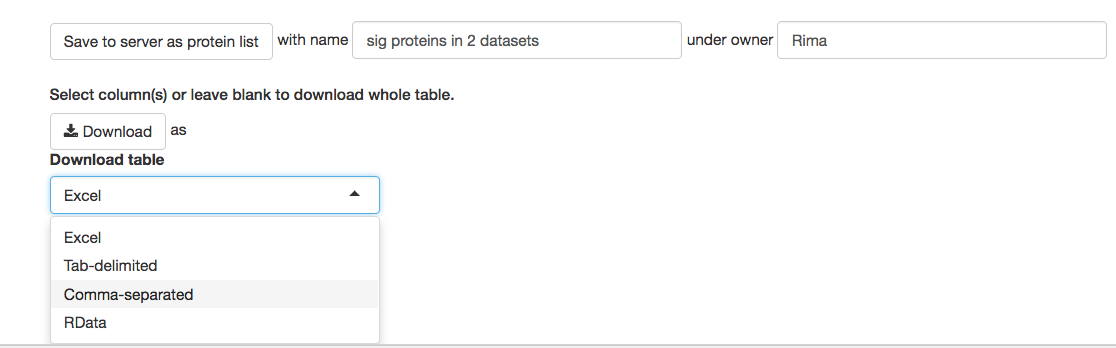

Fig. T11-D
